# GraphVelo allows for accurate inference of multimodal velocities and molecular mechanisms for single cells

**DOI:** 10.1101/2024.12.03.626638

**Authors:** Yuhao Chen, Yan Zhang, Jiaqi Gan, Ke Ni, Ming Chen, Ivet Bahar, Jianhua Xing

## Abstract

RNA velocities and generalizations emerge as powerful approaches for extracting time-resolved information from high-throughput snapshot single-cell data. Yet, several inherent limitations restrict applying the approaches to genes not suitable for RNA velocity inference due to complex transcriptional dynamics, low expression, or lacking splicing dynamics, or data of non-transcriptomic modality. Here, we present GraphVelo, a graph-based machine learning procedure that uses as input the RNA velocities inferred from existing methods and infers velocity vectors lying in the tangent space of the low-dimensional manifold formed by the single cell data. GraphVelo preserves vector magnitude and direction information during transformations across different data representations. Tests on synthetic and experimental single-cell data including viral-host interactome, multi-omics, and spatial genomics datasets demonstrate that GraphVelo, together with downstream generalized dynamo analyses, extends RNA velocities to multi-modal data and reveals quantitative nonlinear regulation relations between genes, virus and host cells, and different layers of gene regulation.

## Introduction

Cells need to constantly detect and adapt to changes in extracellular and intracellular environments. Regulation of their gene transcription is a common mechanism of response. Multiple factors affect the transcriptional activity of eukaryotic genes, including *cis* and *trans* regulatory elements and chromatin structure. High throughput single-cell sequencing data provide the landscape of cell genotype. These data lack, however, information on how the cell state changes over time. Continuous efforts have been made to extract information about gene regulation and develop methods for connecting the cell states to temporal sequences of events captured by single-cell snapshot data. One group of methods that has received extensive attention is based on RNA velocity^1^ for predicting the changes in RNA expression states in the cell. The original RNA velocity method leverages the ratio between nascent and mature transcripts to estimate the rate of change in gene expression. This seminal study has inspired numerous methods for improved RNA velocity estimation based on information from splicing^2–6^, metabolic labeling^7,8^, lineage tracing^9^, and transcriptional factor binding^10^, etc.

The RNA velocity framework has, however, its inherent limitations. First, none of the RNA velocity estimation methods could be applied to any single cell transcriptomic data without restrictions. For example, the splicing-based method is not applicable to prokaryotes or viruses, or organisms without introns. Erroneous inferences of RNA velocities have also been noticed for genes having complex splicing dynamics^11^. Furthermore, it is difficult to estimate the RNA velocities of genes with low expression, which excludes most transcription factors. Second, multi-omics sequencing technologies provide multifaceted information on cellular states alongside the transcriptome modality, and there exist limited systematic methods to extend such velocity estimations to other modalities^12,13^.

The single cell transcriptome state is usually defined by the instantaneous distribution of RNA levels, represented by a multidimensional vector of RNA levels for all (measurable) genes. The usual practice is to map such single cell data onto a reduced space, e.g., transform from a principal component (PC) space to a UMAP representation to facilitate the visualization of the time-evolution of cell state, with each cell state being represented by a point in that space. Numerous dimension reduction and manifold learning algorithms have been developed for representation transformation. In comparison, transforming a velocity vector between representations is a nontrivial task not rigorously addressed in the single cell field. Even worse, a visually correct vector field does not necessarily imply accurate high-dimensional velocity estimation^14^. La Manno et al. proposed a cosine kernel method to address this challenge, which has been adopted since then in most subsequent studies^1^. Li et al. mathematically proved that the cosine kernel asymptotically gives the correct direction of a velocity vector in the large sampling limit, but the magnitude information is completely lost due to a normalization procedure^15^. This loss of information casts concerns when such quantitative information is needed.

In this study, we tackle the above challenges through a graph-theoretical representation of RNA velocities, called GraphVelo, with dynamical systems underpinnings. GraphVelo takes an ansatz that the measured single cell expression profiles and inferred RNA velocities collectively reflect a dynamical process and are connected through a set of dynamical equations. It exploits such additional constraints that couple a high dimensional velocity field and a corresponding single cell state manifold, and enables the generalization of the approach in the context of multi-modal single cell data. While the combined expression and velocity information has been widely used to infer cell state transition trajectories^2,16^, GraphVelo presents the advantage of enabling downstream analyses such as that performed by dynamo^7^ to extract quantitative information on causal gene-gene relations that dictate the cell state transitions. Benchmarking of the proposed graph framework against simulated and experimental single cell data lends support to its broad utility.

## Results

### GraphVelo infers manifold-consistent single cell velocity vectors through tangent space projection and transforms between representations through local linear embedding

Consider that the internal state of a cell can be specified by an *N*-dimensional state vector ***x***, with *N* >>1 generally. Assume that the temporal evolution of the cell state follows a continuous and smooth curve ***x***(*t*) (see Methods for more general mathematical formulation). The instant velocity vector ***v***(***x***, *t*) = d***x***(*t*)/d*t* is always tangent to the curve of ***x***(*t*) (as a function of *t*) at ***x***. One can generalize to the situation that the trajectories of a swarm of cells form a *M*-dimensional manifold ℳ(***x***) embedded in the *N*-dimensional state space with *M* << *N* typically, as revealed by high-throughput single cell omics data. Then under the ansatz that a velocity vector ***v***(***x***, *t*) dictates the evolution of a state vector ***x***(*t*), ***v*** must lie in the tangent space of ℳ(***x***), denoted as *T*_*p*_ ℳ. In practice, the RNA velocity vectors inferred from existing methods do not automatically satisfy this tangent space requirement (but see^4^).

Taking various inferred single cell RNA velocity vectors, e.g. splicing-based, metabolic labeling-based, or lineage tracing-based, as input, GraphVelo takes advantage of the nature of the low-dimensional cell state manifold to: 1) refine the estimated RNA velocity to satisfy the tangent space requirement; 2) infer the velocities of non-transcriptomic modalities using RNA velocities. GraphVelo thus serves as a plugin that can be seamlessly integrated into existing RNA velocity analysis pipelines, and help process single cell data for downstream cellular dynamics analyses using methods such as dynamo (Fig. 1a).

**Figure 1.**
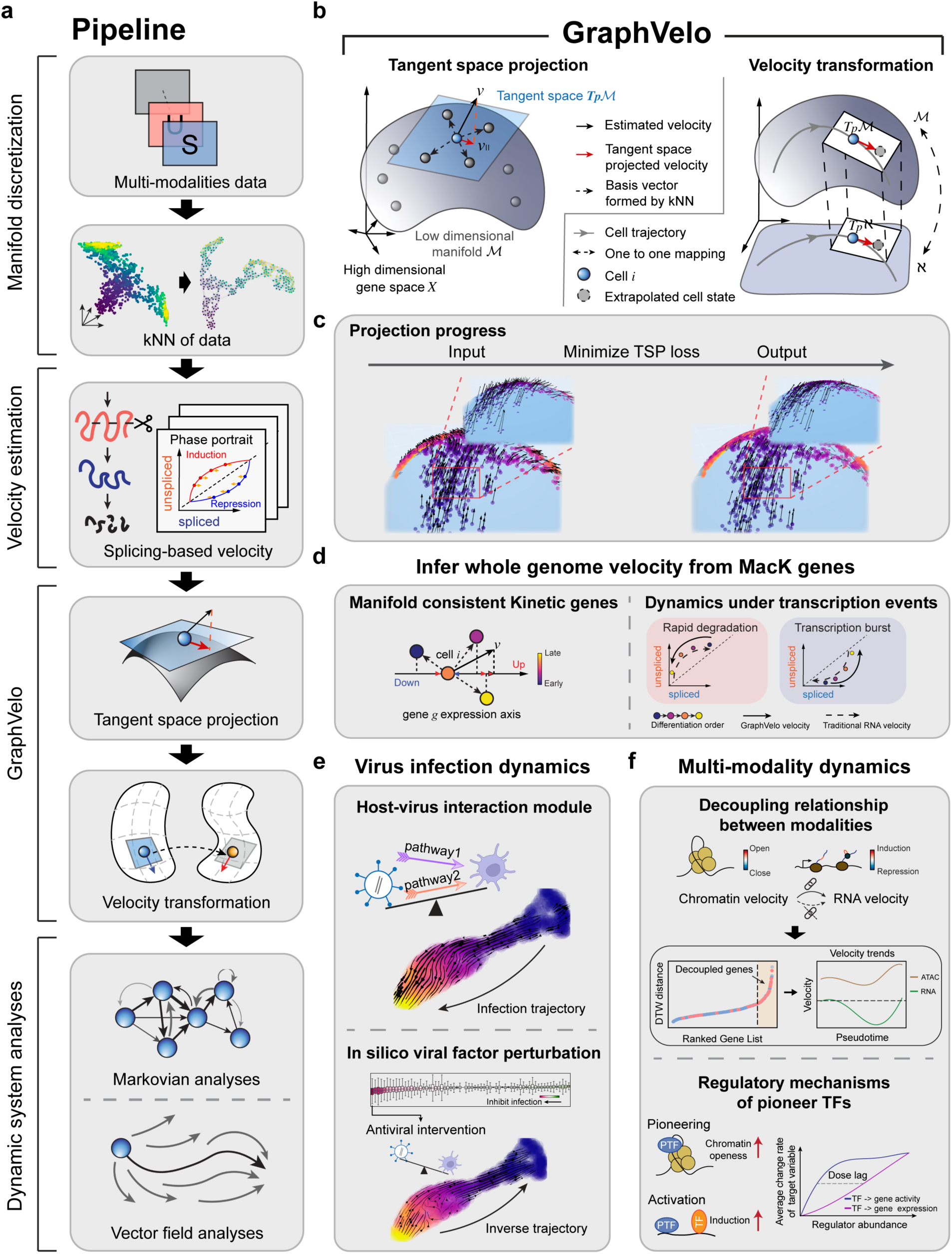
Refining RNA velocity by tangent space projection and transforming between representations using GraphVelo. **(a)** Workflow of RNA velocity-based analyses incoporating GraphVelo. Note GraphVelo takes any form of RNA velocity (i.e., not just splicing-based velocity) as input, and the kNN neighborhood is defined in the full state space (e.g., by both scRNAseq and scATACseq in multi-omics data). **(b)** Schematic of tangent space projection and velocity transformation between homeomophic manifolds. *Left*: RNA velocity vectors are projected onto the tangent space defined by the discretized local manifold of neighborhood cell samples. *Right*: GraphVelo allows for transformation of velocity vectors from a manifold embeded in a higher dimensional space (***𝐾***) to that in a lower dimensional space (ℵ), and *vice versa*. **(c)** The process of minimizing the loss function of tangent space projection. Noisy velocity vectors (*left*) generated by adding random components orthogonal to those sampled from an analytical 2D manifold were projected back onto the 2D manifold, resulting in smooth velocity vectors that lie in the tangent space (*right*). **(d)** GraphVelo allows whole genome velocity inference based on the robustly estimated MacK genes (see also Figure 3). Velocities of genes undergoing variable kinetic rates, such as rapid degradation or transcription burst, are difficult to be correctly inferred by other methods, but can be inferred robustly with GraphVelo. **(e)** Virus infection dynamics and underlying host-virus interaction mechanisms uncovered by GraphVelo (see also Figure 4). *Upper*: pathways involved in host-virus interactions were identified using GraphVelo. *Lower*: GraphVelo predicted reversed trajectory of viral infection in response to *in silico* perturbations of viral factors. **(f)** GraphVelo provides a consistent view of epigenetic and transcription dynamics (see also Figure 5). *Upper*: GraphVelo analyses on multio-mics data revealed that most cell-cycle dependent genes showed decoupling between transcription dynamics and chromatin accessibility change dynamics. *Lower*: Effective dose-response curves reconstructed from multi-omics data revealed pioneer transcription factors increased chromatin accessibility then transcription of targe genes.

Practically, GraphVelo approximates the tangent space at a cell state ***x*** by a *k*-nearest neighbor (kNN) graph following the local linear embedding algorithm^17^, and uses the more reliable data manifold ℳ to refine the velocity vectors by imposing the constraint that the *N*-dimensional velocity vector ***v*** should lie in the tangent space (Fig. 1b&c). Consider a given point ***x***_*i*_on a manifold corresponding to the expression state *i* of the single cell. Its infinitesimal neighborhood forms an Euclidean space that approximates the tangent space *T*_*p*_ℳ. With a sufficient sampling of the neighboring cell states *j* in the state space, the incremental displacement vectors between cell state *i* and its neighboring cell states, ***δ***_*ij*_ = ***x***_j_ − ***x***_*i*_, form a set of complete albeit possibly redundant and nonorthogonal/non-normalized basis vectors of the Euclidean space in the local region. Then the projection of the measured velocity vector onto *T*_*p*_ℳ can be expressed as a linear combination (see also Supplemental Text I),

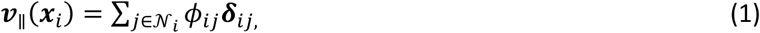

where 𝒩_*i*_ is the neighborhood of cell state *i*, defined by its *k* nearest neighbors in the feature space determined by sequencing profiles. Direct application of eq. 1 to determine the coefficients *ϔ_ij_* is numerically unstable in real data (see Supplemental Text II for detailed discussion). Instead, we performed the projection by optimizing the following tangent space projection (TSP) loss function (Fig. 1c),

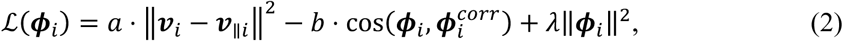

where ‖·‖ refers to vector modulus. 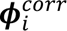 is a heuristic “cosine kernel” widely used in the RNA velocity analyses for projecting velocity vectors onto a reduced space, the elements of which are *ɸ_ij_*(*with j* ∈ 𝒩_*i*_)(see also Supplemental Text III); the second term cos(⋅,⋅) denotes the cosine similarity. The first term in the loss function learns the correct velocity magnitudes, and the second term retains the reliable direction information based on previous mathematical analyses showing that 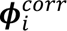 asymptotically gives correct direction of the velocity vector^15^. The L2 regularization is used to bound parameters *ɸ*_*i*_. Hyperparameters *a*, *b* and *λ* are for retaining the projection strength, direction, and for regularization, respectively.

With local linear embedding, it is straightforward to transform velocity between different representations. Assuming a mapping function *f* exists connecting manifold ℳ and ℵ such that for cell *i* with state vector ***x***_*i*_in ℳ, the coordinate of the same cell in ℵ is given as ***y***_*i*_ = *f*(***x***_*i*_). Since a given local patch of continuous manifold is approximated by an Euclidean space, a locally linear transformation connects the patch in the two representations. Consequently, for a vector described by eq. 1 in ℳ , the velocity vector in ℵ is,

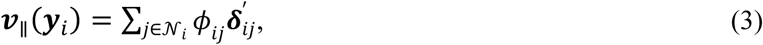

where ***δ***^′^_ij_= ***y***_j_ − ***y***_*i*_. That is, one only needs to change the basis vectors.

Therefore, eq. 1-3 form the mathematical and computational foundation of GraphVelo. With eq. 3 one can extend velocity inference to datasets that velocity inference is not traditionally applicable such as host-virus interactome and multi-omics datasets based on the Whitney embedding theorem^18^. Details of the mathematical foundation were given in Methods. With velocity vectors refined with GraphVelo, one can readily perform downstream analyses, as exemplified in Fig.1d-f, which will be further elaborated below in the context of specific applications.

### Benchmark studies demonstrate effectiveness of GraphVelo across simulation datasets with diverse topology

To demonstrate the effectiveness of the geometry-constrained projection, we first benchmarked our method on a 3D bifurcation system constrained on a 2D manifold (Methods). We added random components vertical to the tangent plane to mimic the noise. The resulting velocity vectors inferred by GraphVelo through minimizing the TSP loss were consistent with the ground truth vectors (Fig. 2a). Both GraphVelo and cosine kernel successfully removed the normal components (Fig. 2b i) and maintained the directional information (Fig. 2b ii), but only GraphVelo kept the velocity magnitude information (Fig. 2b iii).

**Figure 2.**
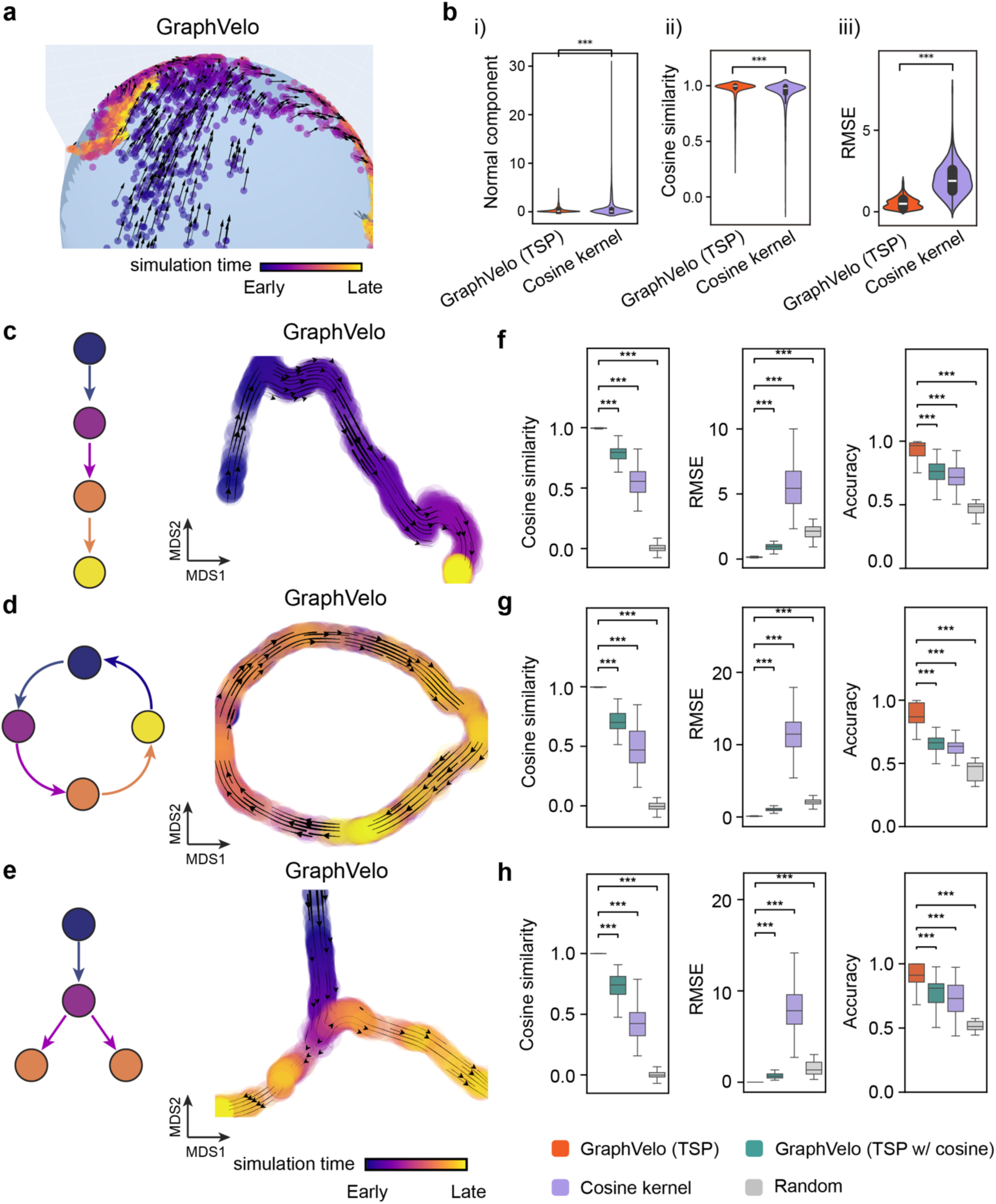
Testing graphVelo on simulated datasets. **(a)** Velocity vectors of an analytical three variables bifurcating vector field constrained to a spherical surface. The data points were colored by simulation time. **(b)** Violinplots of: i) normal component of velocity vectors, ii) cosine similarity and iii) root mean square error (RMSE) between ground truth and velocity vectors projected by GraphVelo and cosine kernel, respectively. **(c-e)** Simulation of scRNA-seq data using dyngen under linear, cycling, and bifurcating differentiation models (*left*), and velocity fields projected on multidimensional scaling (MDS) coodinates (*right*) using GraphVelo-corrected velocities, respectively. Each simulation consists of 1,000 cell states and 100 genes. The cells in different states were colored by their simulation time along trajectory. **(f-h)** Boxplots of cosine similarity, , and accuracy between the ground truth velocity vectors and dyngen simulated velocities after projection using GraphVelo TSP loss without cosine regularization, GraphVelo TSP loss with cosine regularization, cosine kernel, and random predictor, respectively.

Next, we performed multifaceted evaluations of the ability of GraphVelo to robustly recover the transcriptional dynamics across a range of simulated datasets with different underlining phenotypic structures. We used dyngen^19^, a multi-modal scRNA-seq simulation engine, to generate gene-wise dynamics defined by gold-standard transcriptional regulatory networks (Methods). We generated simulated scRNA-seq data for networks with a variety of underlying linear, cyclic, and bifurcating topological structures, and recovered the corresponding vector field using GraphVelo-corrected velocity vectors (Fig. 2c-e). To comprehensively assess the outcome, we used three diverse metrics, cosine similarity, root-mean-square error (RMSE), and accuracy, which evaluate the correctness of velocity direction, magnitude, and sign, respectively. We presented (Fig. 2c-e) the comparative results obtained with the cosine kernel and with GraphVelo (TSP with (i.e., eq. 2 with *b* ≠ 0) and without (eq. 2 with *b* = 0) the cosine regularization term). By minimizing TSP loss, GraphVelo preserved both the direction and magnitude of the vector field (Fig. 2f-h). With an increase of noise level by adding Gaussian noise to the ground truth vectors, GraphVelo refined the distorted velocity and outperformed the cosine kernel projection consistently (Extended Data Fig. 1a-c).

Then, we tested whether manifold constraints could preserve the speed of the cell progression across different representations. GraphVelo was able to scale velocity vectors between the original space and the PCA space, showing a high correlation with the ground truth, even as noise levels increased, whereas the cosine kernel failed (Extended Data Fig. 1d). The results on UMAP showed less agreement, which is not surprising. UMAP is a convenient representation for visualizing single cell data but is not designed for representing quantitative cell state transition dynamics. That is, UMAP is not a continuous transformation from the original gene space and cannot preserve local distances after projection.

To further explore whether GraphVelo could correct the RNA velocity estimated by the splicing kinetics, we took the velocity inferred using different packages (scVelo^2^, dynamo^7^ and VeloVI^5^) as input. The output from GraphVelo agreed significantly better with the ground truth compared to the raw input (Extended Data Fig. 1e), highlighting the significant improvement achieved by GraphVelo in evaluating both the direction and magnitude of the velocity vector fields across all datasets.

### GraphVelo achieved consistent improvements across multiple real-world datasets

Although GraphVelo is designed as a velocity-correction and prediction model complementing existing velocity estimation tools, we benchmarked the performance of scVelo-based GraphVelo outputs on five independent datasets (FUCCI^20^, pancreatic endocrinogenesis^21^, dentate gyrus^1^, intestinal organoid^22^ and hematopoiesis^7^) against five RNA velocity estimation methods (scVelo^2^, VeloVI^5^, UniTVelo^23^, DeepVelo^24^ and CellDancer^3^) (Extended Data Fig. 2-4). GraphVelo achieved noticeably improved cross-boundary correctness (CBC) score^25^ against input velocity and other advanced methods (see Methods for calculation details). We evaluated velocity consistency^2^ across two datasets whose trajectories were estimated using different tools (Extended Data Fig. 4). While GraphVelo shows slightly lower overall vector smoothness scores compared to others, we observed that these models tend to produce overly smooth and homogenized velocity fields, which may obscure biologically meaningful heterogeneity. In contrast, GraphVelo preserves fine-grained local transitions and reveals subtle divergence in vector field, particularly around fate bifurcations in endocrinogenesis data.

We examined the cell cycle datasets annotated by the dynamo package, which features a relatively simple geometry, to quantitatively evaluate our method. First, we focused on the CBC score between cell cycle states. The velocity vectors processed by TSP showed greater consistency with the ground truth compared to the unprocessed inputs (Extended Data Fig. 5a). To demonstrate the effectiveness of GraphVelo in scaling velocities based on the data manifold, we used the L2 norm of velocity vectors to quantify the cell cycle speed. This analysis revealed a peak in velocity within the M and G1 phases, which was also reflected in the distribution of total UMI counts (Extended Data Fig. 5b). We further validated these findings using the cycling A549 cell line sequenced by sci-fate^26^ (Extended Data Fig. 5c) and through the temporal variation of stratified cell cycle speed based on velocities inferred from metabolic-labeling data (Extended Data Fig. 5d). Leveraging the quantitative velocity vectors generated by GraphVelo, we classified genes by both the phase and peak magnitude of their velocities (Extended Data Fig. 5e). Analysis of phase-magnitude relationships uncovered the sequential activation cascade of marker genes throughout the cell cycle.

### GraphVelo infers quantitative genome-wide RNA velocity from a subset of genes with manifold-consistent RNA turnover kinetics

Most RNA velocity methods are based on biophysical models of mRNA turnover dynamics with specific assumptions that may break down in certain cases^11^. These methods typically provide velocities of only a subset of (∼ 500 or less) genes, termed velocity genes in the subsequent discussions, out of a larger list of (∼2-3 k) highly variable genes in a dataset, and some of the velocities are questionable. For example, the splicing-based RNA velocity may have an erroneous sign for processes under active regulation on mRNA degradation or promotors switching between states with different transcription efficiency (Fig. 3a, Extended Data Fig. 6).

**Figure 3.**
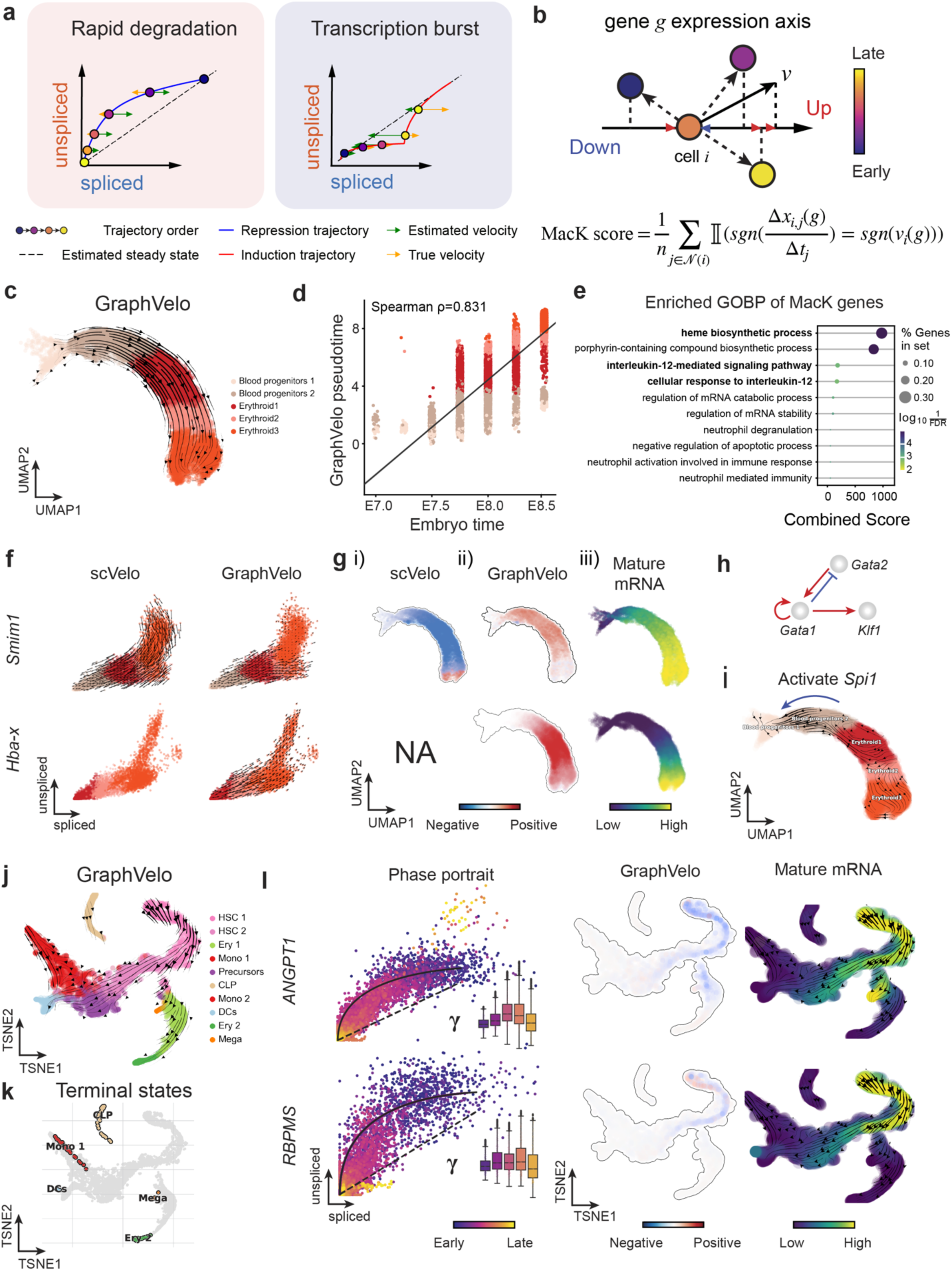
Delineating transcriptome-wise progression with manifold-consistent kinetic genes using GraphVelo. **(a)** Schematitc of transcritional events mislead RNA velocity estimation in the phase portrait by standard approaches. *Left*: for genes exhibiting rapid degradation, the cells appear above the steady state line on the phase portrait, whereas the true velocity is negative. *Right*: For genes exhibiting transcription burst, the transcription rate abruptly increases at intermediate states, leading to a steady state line whose slope is overestimated. **(b)** Schematic of manifold-consistent score calculation for robustly estimated velocity genes. **(c)** The projected velocity field from GraphVelo are consistent with the erythroid differentiation by using all highly variable genes. **(d)** The correlation between GraphVelo vector field-based pseudotime and embryo time for erythroid lineage cells. Spearman correlation coefficients are shown. **(e)** GO enrichment analyses of top ranked MacK genes. **(f)** The phase portrait of two transcription burst genes (*Smim1*, *Hba-x*). **(g)** Scatter plots of: i) velocities estimated by scVelo, ii) refined velocities by GraphVelo, and iii) mature mRNA expression of transcription burst genes (*Smim1*, *Hba-x*). Cells were colored by corresponding velocity, and mature mRNA abundance, respectively, and visualized on the UMAP representation. **(h)** Gene regulatory cascade unraveled by GraphVelo-based vector field analyses that drives cell lineage commitment. **(i)** Activation of *Gata1* inhibitor TF *Spi1* lead to reversed velocity flows in gastrulation erythroid maturation investigated through in silico perturbation analyses on GraphVelo-based vector field. **(j)** Velocities derived from GraphVelo for the branching lineage in the hematopoiesis development and projected onto a pre-defined TSNE embedding. Directions of the projected cell velocities on TSNE are in agreement with the reported differentiation directions. **(k)** Terminal states identified by CellRank based on Markov chain formulation derived from GraphVelo velocities. **(l)** Phase portrait, velocity estimated by scVelo, refined velocity by GraphVelo, and gene expression of mature mRNA of identified rapid degradation genes (*NPR3, ANGPT1*). The cells were colored by the palantir pseudotime^31^ in the phase portrait. The box plots showed cell-specific ***γ*** for cells divided into bins according to pseudotime ordering in the phase protrait.

GraphVelo uses the velocities of high-confidence genes obtained from any method as input to infer velocities of other genes. One can use several existing approaches to evaluate the confidence scores of inferred RNA velocity values of genes^7^. Alternatively, we identified a subset of Manifold-consistent Kinetics (MacK) Genes based on their agreement with prior knowledge or additional information acquired from other methods such as lineage tracing (Fig. 3b), We first applied GraphVelo to a mouse erythroid maturation dataset^27^. This study provided a transcriptional landscape of the erythroid lineage with well-documented differentiation trajectory during mouse gastrulation. Previous analyses have shown that the dataset contains genes with multiple rate kinetics, leading to erroneous prediction of the cell state transition direction^27,28^. We selected the top 200 out of 450 velocity genes as MacK genes, representing those with robustly estimated velocities (see Methods for details). The projected vector field in UMAP showed consistency with prior knowledge in developmental biology (Fig. 3c). We then used the corrected RNA velocities for dynamo velocity field analyses. The vector field-based pseudotime accurately predicted the lineage with scRNA-seq data of temporal mouse embryos (Fig. 3d).

Previous studies identified multiple rate kinetics (MURK) genes showing transcription bursts in the middle of erythroid differentiation^28^. For example, two MURK genes, *Smim1* and *Hba-x*, showed complex patterns of phase portrait (Fig. 3f). Consequently, the RNA velocity of *Simi1* inferred with scVelo was negative along a major part of the developmental axis (Fig. 3g i), contradicting the trend of increasing *Simi1* mRNA levels (Fig. 3g iii). For *Hba-x*, scVelo even failed to infer its RNA velocity. On the other hand, GraphVelo inferred velocities and predicted correct kinetic patterns of these genes (Fig. 3g ii). Similar performances have been observed in other MURK genes (Extended Data Fig. 7a). To examine the overall prediction of cell state transitions from transcription burst genes, we projected the MURK genes velocity inferred from GraphVelo and scVelo to the predefined UMAP. The velocities from GraphVelo but not scVelo correctly captured the directional flow of differentiation using only MURK genes (Extended Data Fig. 7b). We further recapitulated gene succession and oscillation magnitudes along the erythroid trajectory and evaluated the phase-magnitude relationships of all highly variable genes (Extended Data Fig. 8a). Compared to non-MURK genes, MURK genes exhibited larger average velocity magnitudes and were predominantly enriched in the late stages of lineage progression. Analysis of genes with larger peak velocity amplitudes identified *Fth1*, *Car2*, and *Hbb-bs* as candidates with altered kinetic parameters. These dynamics patterns were evident in their phase portraits and gene expression trends (Extended Data Fig. 8b&c), consistent with previous reports that *Car2* transcription in erythroid cells is regulated by both the promoter activity and long-range enhancer interactions^29^—such complex regulations lead to its transcription dynamics not well-described by the simple transcription model used in the original splicing -based RNA velocity inference.

Next, we evaluated the quantitative performance of GraphVelo in estimating cell-wise transition speed. Specifically, we estimated the speed of cell state transition using the norm of velocity vector in high-dimensional space and identified the transcriptional surge stage (Extended Data Fig. 7c). We hypothesized that the MURK genes, which exhibited a sudden increase in transcription rate during this stage, were responsible for the sharp acceleration in cell state transition speed. Using dynamo, we estimated the acceleration derived from the GraphVelo vector field and found that the acceleration value, as the derivative of the velocity vector, demonstrated its potential as a predictor for transcription burst genes (Extended Data Fig. 7d).

With the velocity estimation extended to the whole gene space, we were able to perform comprehensive mechanistic analyses on the entire genome spectrum. First, we calculated the MacK score for each gene using the corrected RNA velocities. We hypothesized that a gene with a higher MacK score indicated a better agreement between its RNA velocity vector and the developmental axis, suggesting that the gene served as a potential lineage-driver gene. We ranked genes based on their scores and performed GO biological process enrichment analyses for the top genes. Indeed, the enriched processes were associated with erythropoiesis, including the heme biosynthetic process and interleukin-12-mediated signaling pathway^30^ (Fig. 3e).

Next, we applied dynamo to perform differential geometry analyses of the vector field and mechanistically dissected the activation cascade of erythroid marker gene *Klf1* (Extended Data Fig. 9a&b). Jacobian analyses based on GraphVelo vector field revealed sequential activation of driver transcription factors (TFs) *Gata2*, *Gata1*, and *Klf1* during erythroid lineage differentiation, with

*Gata1* subsequently repressing the expression of *Gata2* (Fig. 3h, Extended Data Fig. 9c)^7^. To further demonstrate the crucial role of transcriptional factor *Gata1* during erythropoiesis, we performed *in silico* genetic perturbation across all cells. Results showed that both inhibiting *Gata1* and upregulating the *Gata1* repressor *Spi1* lead to a reversal of normal developmental flow (Fig. 3i, Extended Data Fig. 9d). The above analyses collectively suggest that activation of *Gata1* in the blood progenitors biased its differentiation to erythropoiesis, agreeing with experimental reports^28^.

To further evaluate GraphVelo, we tested the method on another dataset of human bone marrow development^31^. This developmental process has complex progressions from hematopoietic stem cells (HSCs) to three distinct branches: erythroid, monocyte, and common lymphoid progenitor (CLP). Again, we used the top 100 out of 454 velocity genes as MacK genes to predict the RNA velocities of 2,000 highly variable genes. The GraphVelo velocity field accurately recovered the fate of cells on the sophisticated transcriptional landscape in contrast to scVelo (Fig. 3j and k, Extended Data Fig. 10a). By combining the likelihood estimated by scVelo with the MacK score, we identified rapid degradation and transcription burst genes whose dynamics deviated from the RNA velocity assumptions (Extended Data Fig. 10b&c). *ANGPT1* and *RBPMS* are two examples which were overall highly expressed in the progenitors and decreased quickly along the trajectories (Fig. 3l), reminiscent of what was shown in Fig. 3a. These genes misled RNA velocity inference with scVelo assuming a constant degradation rate constant. GraphVelo revealed a cell context-specific transcription rate 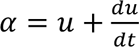 and degradation constant 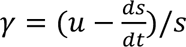 , thus a degradation wave along the differentiation path (Fig. 3l, Extended Data Fig. 10d), consistent with simulation result and reports on regulation of *ANGPT1* mRNA by microRNAs such as *miRNA-153-3p*^32^ (Extended Data Fig. 6d).

It is straightforward to apply Graphvelo to spatial transcriptomics datasets that permit RNA velocity inference. Note that the transcriptional dynamics of a gene can be affected by both the intracellular expression state and extracellular environmental factors. A data manifold containing additional spatial information allows distinction of cell states with similar expression profiles but distinct extracellular environments. Such a refined manifold leads to more accurate inference of the RNA velocities, which are typically performed over averaging the neighborhood of a cell state^1^ (see Methods) and is also used in GraphVelo for tangent space projection. We applied Graphvelo to the mouse coronal hemibrain dataset^33^ processed with a bin size 60, which includes spliced and unspliced transcript information at spatial context (Extended Data Fig. 11a). GraphVelo inferred coherent velocity fields across brain regions, with streamlines on UMAP reflecting anatomically structured transitions that align with the spatial annotation (Extended Data Fig. 11b). Compared to the uncorrected RNA velocities using the dynamo build-in module, GraphVelo captured sharper transcription speed patterns, particularly in the dentate gyrus (DG), a neurogenic region where cell proliferation and neuronal differentiation persist into adulthood^34^ (Extended Data Fig. 11c&d). Spatial mapping of representative genes revealed that GraphVelo velocities were more spatially confined and aligned with known expression domains, whereas the uncorrected velocities were noisier and less localized (Extended Data Fig. 11e). These results highlight the ability of GraphVelo to generate interpretable, spatially structured transcription dynamics in spatial transcriptomic data when splicing information is available.

### GraphVelo reconstructs host–pathogen transcriptome dynamics from infection trajectory

The continuous battle between human immune surveillance and viral immune evasion takes place in the host cell system after viral entry. scRNA-seq data provide a massive and parallel way of assessing the time evolution of both host and viral transcripts, unraveling the delicate inherent dynamics of a virus-host system^35,36^. For splicing-based RNA velocity methods, genes lacking sufficient unspliced transcript counts are typically filtered out—which is inherently the case for viral genes due to their absence of introns. However, several studies^37–39^ have applied tools such as scVelo to host-virus systems by focusing exclusively on host velocity genes that accurately reflect the lineage dynamics, rather than attempting to derive velocities directly from viral transcripts. GraphVelo enables inference of the velocities of virus RNA abundance based on the kinetics of host transcripts velocities, as illustrated next.

We analyzed a human cytomegalovirus (HCMV) viral infection dataset to learn viral transcriptomic kinetics in monocyte-derived dendritic cells (moDCs)^39^. The result from GraphVelo unraveled how viral infection progressed along the transcriptional space (Fig. 4a). The velocity vectors pointed to directions consistent with an increasing trend of the percentage of viral RNAs in individual cells, which inherently served as an indicator of the infection time course^40^. Compared to the trend obtained with the raw RNA velocities from scVelo, the vector field-based pseudotime calculated using GraphVelo-corrected RNA velocities consistently showed higher correlation with the (pseudo)temporal progression of viral infection as reflected by viral RNA percentage (Fig. 4b).

**Figure 4.**
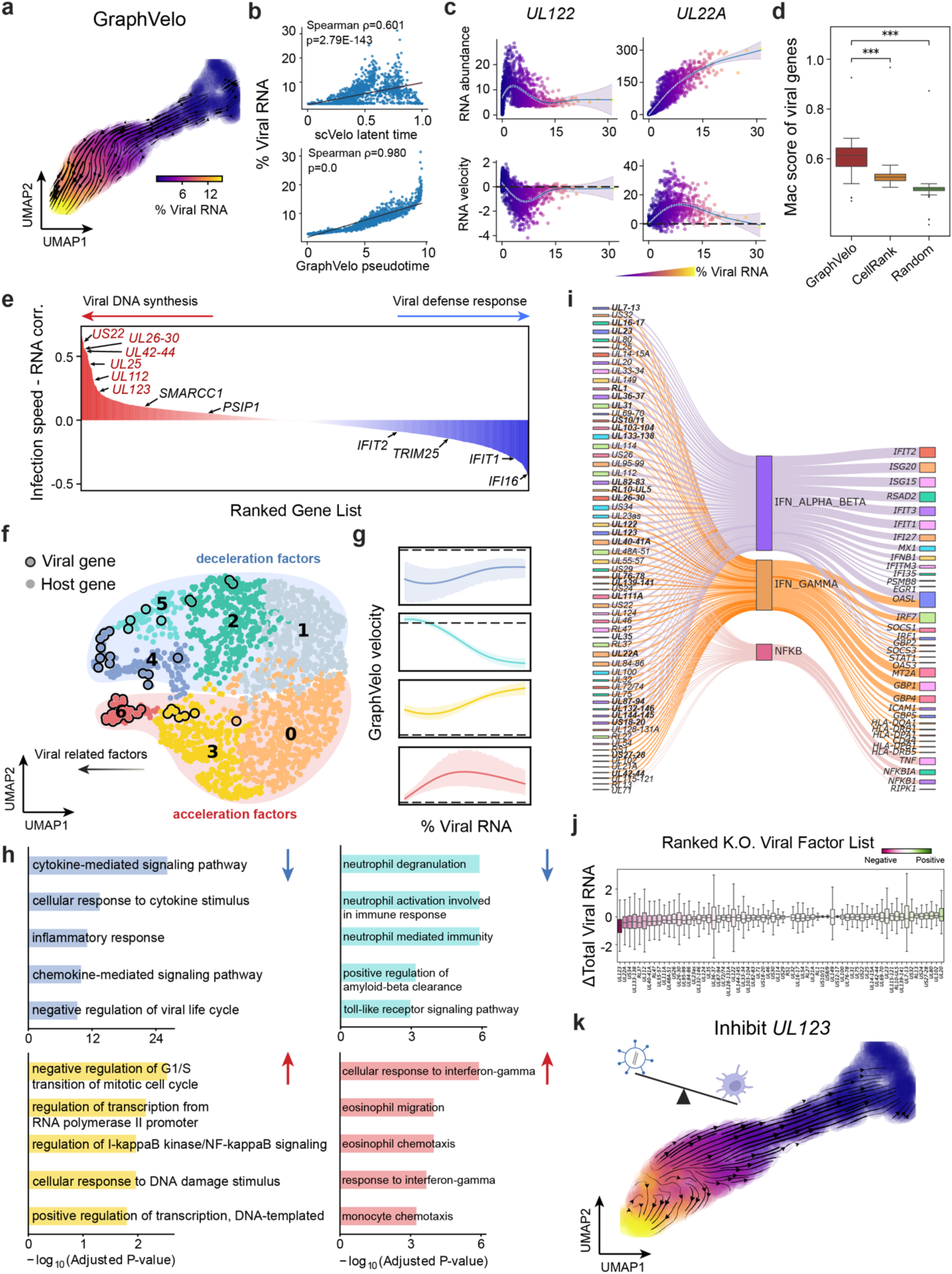
Using GraphVelo velocities to infer host-virus infection trajectory and identify host-pathogen interactions. **(a)** Viral infection captured by the GraphVelo velocity field. Cells were colored by the percentage of viral RNA within a single cell. **(b)** Correlation between viral RNA percentage and pseudotime inferred by scVelo or GraphVelo. Spearman correlation coefficients and *P* values were shown. **(c)** Viral RNA velocities infered by GraphVelo along the viral RNA percentage axis. The black dot line highlights the zero velocity. **(d)** Boxplot summarizing the MacK scores of all viral genes calculated by GraphVelo, CellRank pseudotime kernel and random predictor. **(e)** Correlation between viral infection speed and RNA abundance. Genes were ranked by Spearman correlatioin coefficients. Host and viral genes that contribute to viral DNA synthesis were marked in the left side and those contribute to viral defense response were marked in the right side. Viral genes were highlighted in red. **(f)** UMAP representation of host and viral genes with distances defined by their dynamic expression patterns along the viral RNA percentage axis. **(g)** Example dynamic expression patterns within specific clusters (Leiden4, 5, 3, 6 from top to bottom) along the viral RNA percentage axis. Zero velocity was highlighted by black dot line. **(h)** GO enrichment of each cluster in (g). **(i)** Top host genes inhibited by each viral factor based on dynamo Jacobian analyses. Host effectors were organized by their involved pathways. **(j)** Dynamo prediction of total viral RNA change in response to in silico viral factor knockout. Viral factors were ranked by the mean of total viral RNA changes. **(k)** Vector field change resultant from infinitesmal inhibition of *UL123* during the viral infection process.

Furthermore, the examination of individual genes revealed that the RNA velocities consistently predicted the trend of the mRNA expression level change with increasing virus load (Fig. 4c, Extended Data Fig. 12a&b). Most viral genes started with a fast-increase phase, and the expressions of some genes (e.g. *UL22A*) gradually saturated at high virus load, together with the corresponding RNA velocities approaching zero. One exception is *UL122*, whose expression profile increased first then decreased to a steady state level lower than the peak value. This overshooting is characteristic of a negative feedback network structure^41^. Indeed, a recent study reported that *UL122* negatively regulates its own promotor^42^. Furthermore, comparison of MacK scores across GraphVelo, the CellRank pseudotime kernel, and randomized prediction showed that GraphVelo-computed viral RNA velocities aligned best with the transcriptome gradient of viral load (Fig. 4d). Note that the MacK score also served as a reliable predictor for dynamics-driving factors, specifically viral genes in this case (Extended Data Fig. 12d).

We further quantified the velocity norm of all viral factors as infection speed, and observed that the transcription of viral factors was significantly restricted initially, then gradually increased along the trajectory (Extended Data Fig. 12e). Interestingly, most of the genes that exhibited positive correlation with the infection speed were related to viral DNA synthesis, while those negatively correlated to the infection speed were engaged in host viral defense response^43^(Fig. 4e, Extended Data Fig. 12f).

### GraphVelo identifies host genomic response modules and predicts host-virus gene interactions

With GraphVelo-inferred RNA velocities, we probed the time evolution of lytic infection and the complex interplay between host and viral functional genomes. By fitting the GraphVelo velocity trends along the viral load axis, we identified genes with similar kinetic patterns (Fig. 4c). Using the smoothened velocity trends to calculate the distances, we clustered the genes into seven major modules and visualized them on the UMAP space (Fig. 4f). Not surprisingly, viral genes were concentrated in several enclosed regions, indicating that they formed distinct functional genomic modules during the lytic cycle^40^. The genes showed two major dynamical features: acceleration and deceleration along the viral load axis (Fig. 4g).

To systematically investigate whether the dichotomy between host kinetics genes and viral genes share similar dynamics, we performed gene functional enrichment analyses of host genes residing close to the viral gene clusters. Genes located in the deceleration part were associated with repressing viral genome replication, which includes negative regulation of viral life cycle and known restriction factors in antiviral responses activated in DCs such as the induction of cytokine and chemokine responses as well as interactions with neutrophils^44,45^. In parallel, the toll-like receptor signaling pathway, required for antiviral defense of the host, was arrested^46^. Neutrophil related processes, which typically cooperate closely with DCs to modulate adaptive immune responses^47^, were suppressed. Therefore, the deceleration part also showed how critical set of host factors were silenced by viral entry to achieve immune evasion.

The acceleration groups, on the other hand, demonstrated how viruses hijacked the host cell endogenous cellular programs for virus replication. Notably, the pathways related to viral genome replication were triggered, promoting DNA replication and transcription, such as negative regulation of G1/S transition of mitotic cell cycle^48^, cellular response to DNA damage stimulus^49^ and regulation of transcription from RNA polymerase II promoter^50^. Cells showed a shift towards a transcriptional signature resembling the G1 phase (Extended Data Fig. 12g), agreeing with previous report on HCMV infection^51^. Along the infection process, antiviral interferon (IFN)-γ response of moDC cells was first activated then suppressed. These results highlighted the organized and antagonistic strategies adopted by both host cells and viruses during their tug-of-war for survival and proliferation.

To investigate the crosstalk between host and viral factors systematically in depth, we performed dynamo Jacobian analyses. We scanned the entire spectrum of viral genome and delineated how the HCMV factors silence IFN and NF*κ*B signaling (Fig. 4i). A large proportion of the identified viral factors functioned in evading host cell immune responses, a finding supported by several recent studies^52–54^. *In silico* virus-directed knock out experiments revealed altered accumulation patterns of viral transcripts (Fig. 4j). Notably, inhibition of *UL123*, which ranked first with the total viral RNA inhibition in our analyses, led to a qualitatively distinct trajectory. These results highlight the multifunctional *UL123* locus in the viral genome as a potential target for antiviral intervention^40,55^. The analyses demonstrated potential usage of GraphVelo-inferred velocities for understanding the interactions between viral and host factors, assessing the effects of perturbations on infection, and designing potential antiviral interventions^40^.

### GraphVelo reaveals an abortively infected cell population in SARS-CoV-2 infection

Although HCMV is renowned for its elaborate transcriptional landscape—characterized by extensive alternative splicing, overlapping transcripts, and diverse isoform expression—the RNA virus SARS-CoV-2 transcriptome reveals an even greater level of complexity within its relatively compact ∼30 kb genome^56^. To characterize the molecular mechanisms of host response which protect cells from productive trajectory, we applied GraphVelo to a SARS-CoV-2 infected Calu-3 cells dataset^57^. To focus on the host-virus interactions and their corresponding fate outcomes, we subset the infected cell population for downstream analyses. The infected cluster M, characterized by interferon production genes, was hypothesized to represent a subpopulation of abortively infected cells^57^, similar to those described in herpesviruses HSV-1^58^ and HCMV^40^. We subsampled the infected cell population for downstream analyses and visualization, where cluster M exhibited distinct connectivity properties compared to the main infected groups (Extended Data Fig. 13a). To confirm that M is one of the terminal states of infection outcomes with such low cell abundance (Extended Data Fig. 13b), we applied GraphVelo to gain the quantitative velocity with dynamo outcomes as input (Extended Data Fig. 13c). Using vector field topology analysis, we classified an initial state with relatively low viral load, two terminal states associated with high apoptosis activity, and a saddle point characterized by high viral load (Extended Data Fig. 13d). Notably, the region corresponding to cluster M was identified as an attractor, confirming it represents an abortively infected cell state with a high death rate^57^.

We further validated the complex lineage commitments of SARS-CoV-2–infected cells using the CellRank framework^16^, which successfully identified three potential terminal states as outcomes of host-virus competition (Extended Data Fig. 13e). Interestingly, we characterized the saddle stage in dynamo as a productive terminal state governed by viral genes (Extended Data Fig. 13f), exhibiting high viral transcription speed (Extended Data Fig. 13g). The pathogen-triggered cell death can be driven for protecting the host or for pathogen dissemination purpose^59^. To investigate the underlying causes for host cell death, we performed gene functional enrichment analyses on the top correlated genes along distinct lineages (Extended Data Fig. 13h). Active programs in the abortive infection lineage were highly enriched for host defense mechanisms against viral invasion, consistent with previous studies^40,58^. In contrast, the drivers of the apoptosis-associated lineage revealed a distinct profile. Notably, we observed enrichment for cellular responses to unfolded proteins, which have been implicated in facilitating pathogen-mediated dissemination of infected cells^60,61^. Additionally, *ERBB2* inhibition has been shown to suppress SARS-CoV-2 replication^62^. These findings support the hypothesis that similar cell death outcomes may arise from fundamentally different host responses. Abortively infected cells appear to promote efficient pathogen clearance, likely through cytokine-mediated immune activation that eliminates both the infected cell and the virus. Conversely, cells undergoing virus-induced apoptosis fail to clear the virus, with cell death instead serving as a mechanism for viral escape, immune evasion, and potential dissemination to deeper tissue layers or the bloodstream.

### GraphVelo permits multi-omics velocity inference and chromatin dynamics analyses

The molecular anatomy during cell development entails multiple layers, and how different layers coordinate to regulate gene expression is a fundamental problem. For example, the anagen hair follicle features distinct lineages branching from a central population of progenitor cells. Ma et al^63^. used SHARE-seq to capture both the transcriptome and the epigenome data simultaneously for the lineage commitment process from transit-amplifying cells (TACs) to the inner root sheath (IRS), cuticle layer, and medulla. Upon robust selection of estimated genes following dynamo criteria (Methods), we further refined the RNA velocities of these genes through tangent space projection and obtained the chromatin open/close dynamics from the corresponding scATAC data using GraphVelo. The resultant vector field in the combined transcriptome-epigenome space proved to reconstruct the correct multilineages differentiation paths during the anagen phase (Fig. 5a).

**Figure 5.**
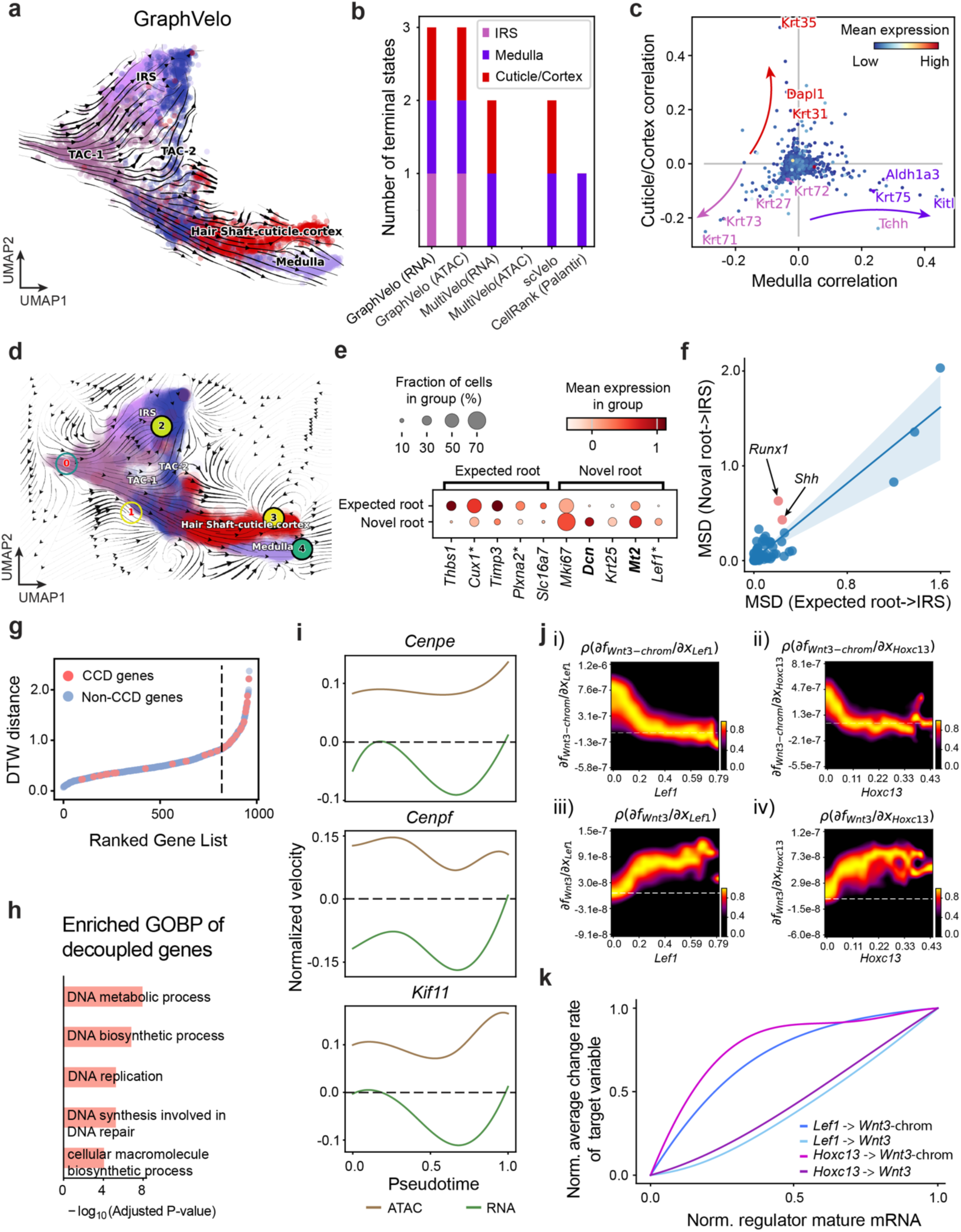
Inferring epigenome and transcriptome consistent dynamics in mouse hair follicle development using GraphVelo multi-omics velocities. **(a)** GraphVelo velocity fields of mouse hair follicle development. Cells were colored by cell macrostates. **(b)** Number of terminal states predicted by CellRank using velocities inferred with different methods. **(c)** Driver genes along multiple lineages identified through CellRank. **(d)** Topological analyses of GraphVelo vector field identified novel root cells and attractors residing in three terminal states(IRS, hair shaft-cuticle cortex, and medulla). **(e)** Expression levels of marker genes in novel root cells and expected root cells. Markers identified by Ma et al.^63^ were highlighted with stars and newly identified markers were highlighted in bold. **(f)** Regression results of MSD values along the transition path from the expected root or novel root to IRS. Two genes *Runx1* and *Shh* genes with large MSD originating from the novel root were highlighted. **(g)** DTW distance between RNA velocity and chromatin velocity of individual genes. CCD genes were colored in red. The dotted line indicates the elbow point separating the decoupled genes from the rest. **(h)** GO enrichment of decoupled genes in (g). **(i)** Line plot of nomarlized RNA and chromatin velocity along pseudotime for genes predicted by GraphVelo to have notable decoupling patterns. Chromatin velocity trends were colored as brown and RNA velocity trends were colored as green. **(j)** Heatmaps of Jacobian element distribution along the axis of regulator RNA abundance of four regulator effector circuits: i) *Lef1* versus *Wnt3* chromatin accessibilities. ii) *Hoxc13* versus *Wnt3* chromatin accessibilities. iii) *Lef1* versus *Wnt3* transcription. iv) *Hoxc13* versus *Wnt3* transcription. **(k)** Effective dose-response curves obtained from integrating the averaged Jacobian elements over the corresponding normalized regulator mature mRNA regulator level in (j).

To test the consistency of dynamics across different modalities, we performed CellRank terminate stage analyses^16^ from the refined velocity vectors. Using GraphVelo velocities of either the RNA modality or the ATAC modality, we accurately estimated three diverse terminal stages (Fig. 5b). For comparison, we also performed similar analyses using MultiVelo, scVelo with all velocity genes or robustly estimated genes in above GraphVelo studies and pseudotime-based vector field inferred by CellRank. The 2D projection of these vector field functions also exhibited seemingly correct velocity flow direction (Extended Data Fig. 14a). However, none of them captured the cell fate commitment based on coarse-grained transition matrix (Fig. 5b, Extended Data Fig. 14b). Notably, the results from the RNA modality and the ATAC modality of MultiVelo gave inconsistent results. GraphVelo-corrected velocities, on the other hand, helped identify the top-correlating genes towards individual terminal populations which showed agreement with previous study^64^ (Fig. 5c, Extended Data Fig. 15).

Next, we conducted differential geometry analyses based on the composite GraphVelo vector field. We identified novel root cells, which were also characterized by chromatin potential (Fig. 5d)^63^. These novel root cells expressed distinct marker genes compared to the expected root cells using the Wilcoxon test (Fig. 5e). Moreover, we unraveled differentially expressed markers identified by the original study^63^, as well as new differentiation-potent genes and validated their initiation properties in another transcriptome dataset (Extended Data Fig. 16a)^64^. To further investigate how these two distinct groups of root cells convert to other cell types, we performed a least action path (LAP) analysis between different cell phenotypes. The expected and novel root cells converted to the IRS terminal state following two distinct least action paths in the vector field (Extended Data Fig. 16b&c). The two paths revealed different temporal change patterns of transcription factor expression profiles (Extended Data Fig. 16d). We calculated the mean-squared displacement (MSD) for every transcription factor to explore the dynamics of TFs along the path from novel root to IRS. The result demonstrated that the fate conversion by novel root was mediated by the *Shh-Runx1* signaling axis (Fig. 5f, Extended Data Fig. 16d), which has been demonstrated in human embryonic stem cells^65^ and is crucial for hair development^66^. In summary, GraphVelo unraveled the multiple molecular mechanisms that orchestrated hair follicle morphogenesis.

With available chromatin velocity and RNA velocity, we set to quantify the coupling/decoupling relationships between chromatin structure and gene expression for each gene (see Methods). Here chromatin structure refers to the extent of exposure/accessibility of the gene locus to the environment as indicated by scATACseq data, shortly called open or closed state; and chromatin velocities refer to changes between these states as inferred from scATACseq counts at the examined gene locus. We used dynamic time warping (DTW) distances between the velocities from different omics layers to quantify the similarity between temporal patterns of these two modalities for each gene. A higher DTW value indicates higher similarity. Using the elbow of the ranked distance curve as a cutoff we identified genes that showed decoupled transcription and chromatin structure dynamics. These decoupled genes had an accumulation of cell cycle-dependent (CCD) genes found in previous study^20^ (Fig. 5g). This group of genes showed strong involvement in cell cycle-related processes, as indicated by GO enrichment analyses (Fig. 5h). Close examinations indicated that the transcription of cell cycle related genes decreased along the differentiation path, while the chromatin structure at the corresponding loci remained open (Fig. 5i, Extended Data Fig. 17). To validate this hypothesis, we further applied GraphVelo to a recently published 10x Multiome dataset from developing human cortex^67^ (Extended Data Fig. 18a). Following the same analyses, we identified decoupled genes and found out that most of these genes were related to cell cycle (Extended Data Fig. 18b-f). which has also been reported in a previous MultiVelo study^13^.

We further performed dynamo differential geometry analyses on the composite transcriptome-chromatin vector field. One intriguing phenomenon observed in lineage dynamics is that *Lef1* and *Hoxc13* are the driver TFs correlated with domains of regulatory chromatin (DORCs) of *Wnt3* ^63^. Differential geometry analyses on the composite vector field can go beyond correlation analyses and provide an underlying casual mechanism. As a prerequisite for such analyses, GraphVelo-inferred RNA and chromatin velocities of the three genes correctly predicted the trend of change of mRNA and ATAC-seq counts (Extended Data Fig. 19a&b), in contrast to the performance of MultiVelo (Extended Data Fig. 19c). Then, Jacobian analyses on the GraphVelo vector field confirmed that priming activation of *Lef1* subsequently activated the *Hoxc13* TF^68^ (Extended Data Fig. 19d&e). Both *Lef1* and *Hoxc13* were found to activate the *Wnt3* target gene, initiating lineage commitment (Extended Data Fig. 19f &g). To quantitatively understand how the two TFs affect *Wnt3* chromatin structure and transcription, we plotted the response heatmap to reflect the distributions of Jacobian elements versus the abundance of mature mRNA for each TF (Fig. 5j). The two terms 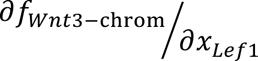 and 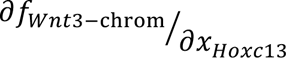 started with positive values at low concentrations of TF mRNA copy numbers then decreased to zero, indicating that increasing the level of either TF lead to further opening of the *Wnt3* chromatin region, and the effect saturated at high TF expression. The other two term 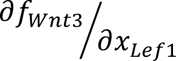 and 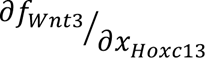 increased with the TF levels, indicating that these two TFs also activated *Wnt3* transcription. Upon integration of the Jacobian elements over regulator expression changes, we obtained the effective dose-response curves obtained (see Methods), which revealed more transparently the TF dose lag between the opening of the target chromatin region and the initialization of transcription (Fig. 5k).

Our results therefore illustrated the sequential events that these two driver TFs, *Lef1 and Hoxc13*, drove as pioneer transcription factors (PTFs) to initiate local chromatin opening and then activated the transcription of *Wnt3*. Notably, computational methods and experimental research confirm that *Lef1* acts as a nucleosome binder and exhibits diverse binding patterns across various cell lines^69^. The *Hox* family of TFs has also been shown to have the capacity to bind their targets in an inaccessible chromatin context and trigger the switch to an accessible state^70^, consistent with our analyses that *Hoxc13* revealing that it shared a regulation mechanism similar to that of PTF *Lef1*.

## Discussion

In this work, we provided a general framework that extends the framework developed for RNA velocity and related approaches to various data modalities such as proteomics, spatial genomics, 3d genome organization, and imaging data, which were originally beyond reach of this framework. We validated GraphVelo using various *in vivo* cellular kinetics models, confirmed its efficacy and robustness in handling complex and noisy multimodal data. Upon application to various datasets, we unraveled gene regulation relations of an extended list of genes, host-virus gene regulations, and coupling between transcription and local chromatin structures. GraphVelo can be seamlessly integrated with broad downstream analyses, such as dynamo continuous vector field analyses, as well as Markovian analyses using graph dynamo or CellRank.

## Methods

### Dynamical systems theory formulation of cellular state transitions

Assume that a cell state can be represented by the cell volume *V* (or cellular compartment size) and the copy number of *L >> 1* pairs of gene products (***n****_m_*, ***n****_p_*), where *m* and *p* designate mRNA and protein, and the bold fonts indicate vectors. For simplicity here we only consider *m* and *p*. It is straightforward to generalize to finer cell state specifications, for example, with distinction of nuclear and cytosol localizations, posttranslational states of proteins, other species such as microRNAs, epigenetic states, etc.

The temporal evolution of the cell state is described by a set of chemical master equations. When the copy numbers of molecular species are not too small, and the chemical reactions are not strongly coupled, Gillespie showed that the chemical master equations can be approximated by a set of chemical Langevin equations^71^. With extrinsic noises also included, we assume the ansatz that the dynamics of cell state is described by a set of generic stochastic differential equations,

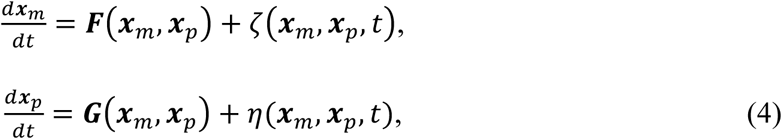

where the *L*-dimensional vectors ***x***_*m*_ and ***x***_*p*_ are cellular concentrations of *m* and *p*, and the *ζ* and *η* are taken as white noises with zero mean.

The low-dimensional manifold assumption is central to machine learning approaches on data analyses. From dynamical systems theory perspective, after a transient time a multi-dimensional dynamical system often converges to a low-dimensional slow manifold. In practice such property has been exploited with techniques such as quasi-steady-state approximation, quasi-equilibrium approximation. For a rigorous formulation, assume that one can identify a set of variables (***z***, ***Z***) with (*2L-M*) dimensional fast variables ***z*** = ***z***(***x***_*m*_, ***x***_*p*_) and *M*-dimensional slower variables ***Z*** = ***Z***(***x***_*m*_, ***x***_*p*_). Xing and Kim extended the celebrated Zwanzig-Mori projection^72,73^ to a general dynamical system described by eq. 4^74^. The projection procedure results in a set of stochastic integral-differential equations of ***Z*** with colored noises, which are formally equivalent to eq. 4. Then if one assumes clear time scale separation between ***z*** and ***Z***, the equations reduce to a set of Langevin equations with white noises, 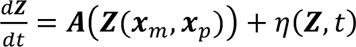, where (***Z***, *t*) are white noises with zero mean. Through ensemble averaging over the vicinity of a given point ***Z***, one has

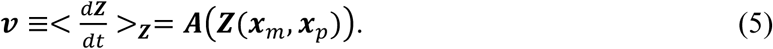

The equations define a *M*-dimensional manifold embedded in the B***x***_*m*_, ***x***_*p*_C space. A scRNA-seq data then measures the corresponding manifold projected to the transcriptomic subspace.

One should note that in practice the reported RNA velocity vector of a cell state *i* is typically obtained through averaging the raw velocity vectors of cell states within its neighborhood 𝒩_*i*_ on the manifold as a numerical approximation of the ensemble average, 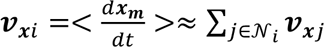. In this work, we used the k-nearest neighbor (kNN) algorithm to define the neighborhood in single modality datasets, including spatial-transcriptomics datasets. For multi-omics data, the neighborhood of a cell state was defined using weighted nearest neighbors (WNN)^75^ in the composite cell state space. The procedure of applying GraphVelo is the same for different data types except using the neighborhoods defined on their corresponding data (cell state) manifold.

### Mathematical foundation on applying GraphVelo to multi-omics datasets

Equation 3 in the main text applies to transformation between a manifold embedded in a state space and in a subspace. According to the Whitney Embedding theorem^18^, any smooth real *M*-dimensional manifold can be embedded in a *2M*-dimensional real space provided that *M* > 0. Consider a full set of genes versus a subset in a scRNAseq dataset, or a combined scRNAseq/scATACseq multi-omics dataset versus the scRNAseq subset. Assume that the full cell state space has a dimensionality *N*, while a single cell data manifold is typically low-dimensional with *M* << *N*. Then the Whitney Embedding theorem^18^ suggests that with proper choice of the subset the manifolds in the full space and the subspace are homeomorphic or at least piece-wise homeomorphic (Fig. 1b, Supplemental Text IV), i.e., a one-to-one mapping exists between the two. Then applying eq. 3 allows one to infer the velocity vectors for the full-space representation from those of the subspace.

#### Denoise velocity vectors in the space of principal components (PCs)

Learning coefficients *ϕ*_*i*_in the gene space directly often fails due to thousands of gene profiles. To avoid the curse of high dimensionality and learn parameters in a compact manifold, we designed a procedure to denoise the velocities in a reduced PCA space. Specifically, we extrapolated the cell state *i* in the original space using the infinitesimal propagation operator to extrapolate the future state:

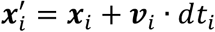

Moreover, we estimated an optimal step size *dt* based on the local density to guarantee the cell states are bound to the manifold:

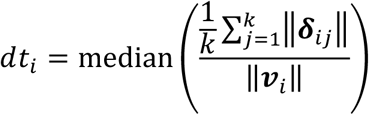

After utilizing the cell-dependent *t*_*i*_to forcing the predictions inhabiting regions of the phenotypical manifold, we applied dimension reduction to project both current and future status from the gene space to the PCA space through linear transformation. Then we obtained the projected velocity vectors as:

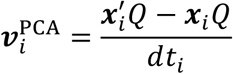

where *Q* is the PC loading matrix estimated using x that serves as the coordinate transformation matrix.

#### Manifold-consistent kinetic genes

The kinetic assumptions between nascent and mature RNA fail when the underlying parameters shift along the developmental trajectory^11^, which leads to transcription burst and rapid degradation in the phase portraits (Fig. 3a). An internal clock exists during cell proliferation and differentiation. Current methods rely on different criteria to select confident estimated velocity genes (see Supplemental Text V for detailed discussion). Here, we presume that the velocity of robustly estimated genes should be consistent with the (pseudo)time derivative estimated under the manifold assumption. We can utilize any available *t*s inferred from data manifold by either pseudotime, velocity latent time, or lineage tracing to approximate the temporal information and use *k* nearest neighbors to define the locally linear plane. After ordering cells within the local Euclidean space, we calculate the MacK score for any gene *g* as an indicator of whether the sign of estimated velocity agrees with the dynamic cascades within manifold,

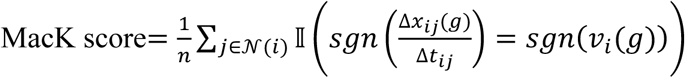

Where 𝒩_*i*_ indicates the neighbor points of cell *i*, 𝕝 represents the indicator function and *sgn* returns the sign of the values. Δ*x_ij_*(*g*), *v*_*i*_(*g*) are the difference in abundance of gene *g* between cell *i* and *j*, and the velocity of gene *g* in cell *i*, respectively. We parallelize the calculation to scale efficiently with the number of genes, which is important due to the number of highly variable genes. We want to point out that one can use methods other than the MacK score to identify genes with reliable velocity estimations.

#### Dynamo criteria

Dynamo^7^ offers a correction strategy by removing genes with low gene-wise confidence in the phase plane. This allows us to identify genes that appear in incorrect phase portrait positions and contribute to erroneous flow directions (illustrated in Fig3. a). To filter out genes with misleading dynamic patterns, one can supply the established lineage hierarchy information to the *dyn.tl.confident_cell_velocities* function in dynamo. This function scores each gene based on the agreement of its behavior in the splicing phase diagram with the input lineage hierarchy priors.

### Post-GraphVelo analyses

#### Reconstruction of extended dynamo vector field from multi-omics data

Dynamo is a general framework of reconstructing dynamical models from scRNAseq data, and it is straightforward to generalize to multi-modal data. The framework is based on specific realizations of eq. 5, 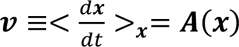, with state vector ***x*** being the transcript concentrations for scRNAseq data, and combined transcript concentrations and continuous quantification of locus-specific chromatin open-close state for multi-omics scRNAseq/scATACseq data. The variables ***x*** can be defined in various representations, e.g., the original gene space, principal component subspace, latent space defined by variational autoencoder, etc. With GraphVelo it is straightforward to transform ***v***_***x***_between different representations.

The continuous vector field functions ***A***(***x***) contain quantitative regulation relations between genes that are learned from single cell data points of (***x***, ***v***_***x***_). Various algorithms can be used to learn the *analytical* forms of ***A***(***x***). The original dynamo paper illustrated a Reproducing Kernel Hilbert Space (RKHS) representation method. The method expresses ***A***(***x***) as a linear combination of pre-selected basis functions, ***v*** = ***A***(***x***) = ∑_***α***_ ***Γ***_α_(***x***), similar to the more familiar Taylor expansion that uses a linear combination of polynomial functions to represent a continuous analytical function. It should be noted that the basis functions and so ***A***(***x***) are generally *nonlinear* functions of ***x***. Following Qiu et al.^7^, we chose Gaussian functions centered at selected reference points *x̃_α_*, 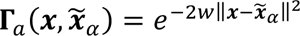, with default parameter value of *w* in the package dynamo. Then we determined the coefficient vectors ***C***_***α***_ through minimizing the loss function 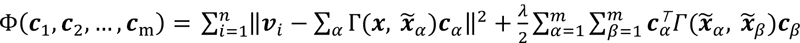, where the first sum was over all the data points, and the second term was Tikhonov regularization weighted by *λ*. The superscript *T* means matrix transpose. One can also use neural networks, e.g., variational autoencoders, to learn an optimal set of basis functions, and other algorithms such as neural ODE to learn ***B***(***x***). The difference is merely algorithmic under the same framework of dynamical systems theories.

##### Jacobian analyses and reconstruction of effective dose-response curves of gene regulation

The extended dynamo vector field contains generally nonlinear relationship about regulation between genes, and between genes and other modalities (e.g., chromatin open/close conformations). Several posterior interpretation methods exist to analyze the vector field. Below we will describe two of them.

With the analytical form of **F**(x), one can calculate efficiently the Jacobian field **J** at any cell state **x**. Each element of **J**, (*J_ij_* = **x**_*i*_⁄*x_j_*), can be understood as an in-silico perturbation experiment on how upregulating gene *j* affects the transcription rate of gene *i,* with all other gene expression levels kept constant at state **x**. For example, a positive value of *J_ij_* indicates that at **x** further increasing the expression of gene *j* causes increase of the transcription rate of gene *i*. Note that the sign and value of a Jacobian element alone does not unambiguously reflect the nature of the regulation. A close-to-zero Jacobian element can be associated with either no regulation or the regulator is at a saturating concentration of regulation. The regulation relation can be direct, or indirect through intermediate molecular species not implicitly treated as variables of the vector field function.

Complementary to the local Jacobian analysis is to reconstruct effective dose-response (D-R) curves of a regulator-target gene pair. The curve reveals the rate of change of one quantity, e.g., the transcription rate or the chromatin open/close dynamics of the target gene, as a function of the value of a regulator, e.g., the mRNA level of a regulator or the chromatin open/close status of a specific genomic region. Note that the D-R curve is generally a multi-variate function, we designed a procedure to reconstruct an effective one-variate function^7^. One can genetically write the regulation on quantity *x*_*i*_ as two terms with and without dependence on variable *x*_j_,

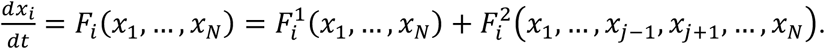

Notice that 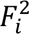 is not a function of gene *j*. First, we calculated the Jacobian element 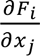 for each measured cell state. Note 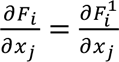, so the background variation from 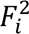 due to effects of other genes has been numerically removed. Then from the histogram of 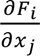 versus *x_j_*, we binned 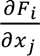 over *x*_j_ and calculated 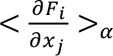, which was averaged over all data points of 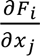 within bin ***α***. Next, we performed numerical integration to obtain 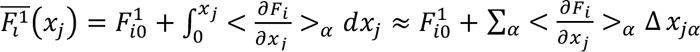. In practice, if 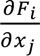 shows large variance within each bin of *x*_j_, it may imply that other factors affect the D-R curve. For example, the regulation of *x*_j_ on *x*_*i*_ may even be opposite at the presence or absence of a specific cofactor. In this case, one should first cluster cells, e.g., grouping cells based on whether an identified cofactor reaches a threshold value, then perform the D-R curve reconstruction on individual clusters.

##### Markovian analyses

Marius et al.^16^ have developed a framework named CellRank to study cellular dynamics based on Markov chain formulation. We use CellRank to identify cell state transitions using velocity kernel and identify terminal states within datasets by GPCCA function module. In addition, CellRank pseudotime kernel^8^ is used for methods comparison in real datasets.

##### Analytical function form of a vector field and differential geometry analyses

Dynamo learns a nonlinear function form of RNA velocity vector field, providing a physics-informed framework which integrates mechanism modeling and single cell data analyses. We use dynamo to learn continuous vector field functions and perform differential geometry analyses such as gene acceleration, vector field-based pseudotime, least action path (LAP), Jacobian analyses and in silico perturbation.

##### Pseudotemporal orderings

GraphVelo itself does not compute an ordering index of cells as we are seeking for a more quantitative method to infer RNA velocity. With an accurate RNA velocity as input, we can approximate the vector field precisely. Thus, we use the scalar potential estimated from the functional form vector field with Hodge decomposition as a proxy of time, which is implemented by dynamo package.

##### Cross boundary correctness scoring

While the GraphVelo framework is designed to quantify velocity vectors across different representations, known transitions between coarse cell states— such as cell types or cell cycle phases—can be used to evaluate the correctness of velocity directions. Suppose there are two cell populations A and B, with A a progenitor state of B. One can define the set of boundary cells between A and B as

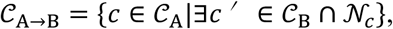

where 𝒞_A_ or 𝒞_B_ denote the sets of cells in state A or state B, 𝒩_*c*_ indicates the kNN of cell *c*. The CBC score is then defined as

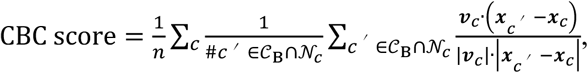

where #*c*′ ∈ 𝒞_3_ ∩ 𝒩_*c*_ is the number of cells in state B which is also the kNN of cell *c*. While the (***x***_*c*_, ***v***_*c*_) can be represented in different basis (raw count, PCA or UMAP), we computed the CBC score in the original count space to ensure that all genes contribute to the velocity estimation. We deliberately avoided using 2D embeddings like UMAP for this purpose, as such visualizations may distort the true geometric relationships in the high-dimensional space and could lead to misleading interpretations.

##### Velocity consistency scoring

For most of the cases, we expect the inferred RNA velocity vectors to be coherent in a uni-directional vector field. To quantify the local consistency of the velocity flow for each cell, we calculate the velocity consistency score^2^ for each cell *i* as the mean correlation of its velocity *v*_*i*_ with velocities from neighboring cells,

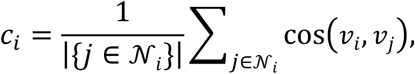

where cell *j* is the neighbor of cell *i* and cos indicates the cosine similarity. One thing should be clarified is that overly smooth and homogenized velocity fields may obscure biologically meaningful heterogeneity.

##### Approximate smooth velocity trends with generalized addictive model

The variation of transcription rates contains the high order dynamic information of the cell system. To model the dynamic patterns of RNA velocity along transition path from noisy data, we refine the velocity vectors by local geometry via TSP and further fit GAM to velocity value of each gene that has been refined by GraphVelo. For any gene g, we model the velocity trend for the temporal variable *t* via

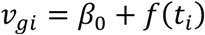

Where *v*_g*i*_ indicates the velocity of gene g in cell *i*, *f* is built using penalized B-splines which allow us to automatically model non-linear mapping while maintaining additivity^76^. To visualize the velocity trends, we select 100 equally spaced testing points along transition path and predict gene expression at each of them using the fitted model. The estimated velocity trends can be treated as smoothed time series for further analyses.

##### Clustering velocity trends along infection trajectory

With manifold-constrained velocity estimated by GraphVelo, we are able to cluster genes into different functional modules which are involved in the same regulatory circuit. We recover transcription variation of both host and virus factors along infection trajectory by fitting GAMs in temporal indicator, the percentage of viral RNA. Next, we select 100 equally spaced time points and generate the GAM-smoothed velocity trends. We compute a kNN graph and cluster the kNN graph using the leiden algorithm. We used k = 15 for the velocity-trend kNN graph and the leiden algorithm with resolution parameter set to 0.3 to avoid over clustering the trends. For each recovered cluster, we compute its mean and standard deviation (pointwise, for all generated points that were used for smoothing) and visualize the smoothed trends per cluster.

##### Characterizing decoupling genes based on dynamic time warping distance between multi-modality velocities

We perform the DTW distance calculation by dtaidistance package. To eliminate the influence of scale in different modality, we maximum-normalize the chromatin/RNA velocity to the same range of [0, 1]. Then we fit the velocity trends of both modalities along vector field-based pseudotime to yield the smoothed velocity trends. We calculate the DTW distance between velocity trends per gene. To distinguish the decoupling genes based on their multi-modality velocities, we rank the genes based on the DTW distance and identify elbow point as cutoff. Genes with distance metric larger than the cutoff are characterized as decoupling genes and used for visualization and functional analysis.

### Synthetic datasets

#### Generating two genes bifurcation process and mapping it to 3d

The bifurcation data (n=2,000 cells) for the toggle-switch system is simulated using the Gillespie algorithm. We use activation and inhibition Hill functions to model the induction and suppression effects between the two genes:

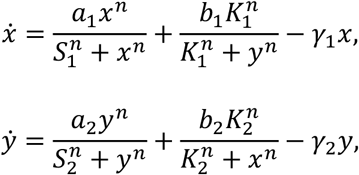

We use the simulation backend implemented by dynamo with default parameters except the timescale (reset *τ* =1) to generate the bifurcating process. We then map the synthetic dataset onto a sphere (radius *r*=70) and yield the variable *z* as:

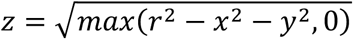

Then we are able to calculate the correctly-scaled 3d vectors by infinitesimal propagation operator with sufficient small step size (*dt*=1 in our case):

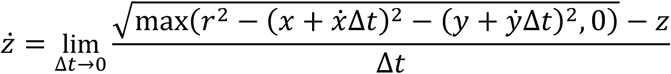

#### Generating scRNA-seq synthetic data with dyngen

To generate high-dimensional single-cell transcriptomic data in silico, we use a multi-modal simulation engine, dyngen, to account for different developmental topologies. We constructe module networks to represent regulatory cascades and feedback loops driving progressive changes in gene expression and influencing cell fate decisions. We generate three datasets with 1,000 cells and 100 genes using the linear, cyclic, and bifurcating loop backbones provided by dyngen, with all other parameters set to default values.

These datasets include simulated nascent and mature mRNA counts along with ground-truth RNA velocities and known manifold structure.

#### Simulation of rapid degradation and transcription burst events on phase portrait

As for genes with variable degradation rate, we present a minimal regulatory network with linear model in which an external signal both inhibits transcription and promotes microRNA (miRNA). The miRNA exerts a linear influence on the degradation rate of mRNA. We have miRNA’s velocity as

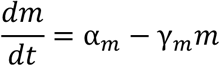

The nascent gene transcription rate and mature mRNA’s degradation rate would change to

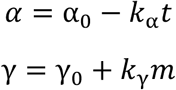

where α_0_ and γ_0_ represent the constant transcription rate and degradation rate without the effect of miRNA, *k*_α_ and *k*γ represent the magnitude of influence from miRNA, *t* indicates the simulation time to mimic the cell-context change along trajectory.

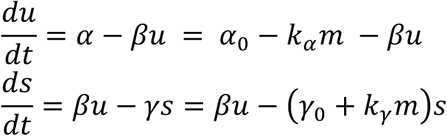

We set the initial condition to a steady state with γ_0_ = ***α***_0_⁄*β* and *s*_0_ = *α*_0_⁄*γ*_0_, while the miRNA abundance *m*_0_ = 0. We simulate *u_t_* and *s_t_* as the microRNA signal *m*_*t*_ gradually increases. The aim is to evaluate whether the estimated RNA velocity consistently aligns in sign with the ground truth RNA velocity.

To generate genes with transcription burst phase portrait, we set the initial condition to γ_0_ = 0, *u*_0_ = 0, together with γ as constant and ***α*** promotes to ***α***^′^ = 3***α*** when simulation reaches specific time.

### Processing of sequencing data

All sequencing data in this study are downloaded publicly (see details in the ‘Data availability’ section). Though the number of confident-estimated gene sets differed case by case, GraphVelo predicted the velocity of all highly variable genes and used them for downstream calculation such as CBC score, CellRank velocity kernel and dynamo vector field learning. We parallelized the TSP optimization to scale efficiently and set the hyperparameters in loss with *a* = 1, *b* = 10 and *λ* = 1 as default in all studies (Extended Data Fig 20).

#### Analyses of scRNA-seq datasets

For the erythroid lineage of the mouse gastrulation, we follow the standard data pre-processing procedures implemented by scVelo and select 9,815 cells and 2,000 highly variable genes to construct the k-nearest neighbor graph using 30 nearest neighbors for downstream calculation.

For the human bone marrow dataset, we follow the standard data pre-processing procedures implemented by scVelo and selected 5,780 cells and 2,000 highly variable genes to construct the *k*-nearest neighbor graph using 30 nearest neighbors for downstream calculation. To estimate the variation of degradation rate *γ* along the differentiation lineages, we divide the cells into five discrete time bins based on precomputed Palantir pseudotime. We then estimate cell-specific degradation rates and visualized their distribution shifts along the hematopoiesis trajectory.

#### Analyses of spatial transcriptomics dataset

For the mouse coronal hemibrain spatial dataset, we follow the monocle preprocessing pipeline implemented in dynamo and select 7,765 cells and 2,000 highly variable genes. To include spatial information during manifold, we built a spatial kNN graph with k=8 and then took the union with the kNN graph based on transcriptomic data. The combined graph is used for downstream analyses.

#### Analyses of HCMV dataset

We sample the cells from donor 1 to eliminate sample-specific variation and further filter out cells lacking immediate early *UL123* gene expression to focus on the viral infection trajectory. We adapt monocle preprocessing recipe implemented by dynamo and yield 1,454 cells and 2,000 highly variable genes for further analyses. The number of nearest neighbors were set to 30 as default. Then 1,022 velocity genes are used for downstream analyses.

We collect the pathway-related genes from MSigDB and perform the Jacobian analyses implemented by dynamo, using viral genes as regulators and host genes as effectors. We rank the regulation relationships based on the collections of Jacobian elements. We pick top 50 inhibited effectors of viral genome and select the common set between pathway genes and all the effectors for visualization.

In silico knock out experiments are performed via dynamo. We suppressed every single virus factors per time using *dynamo.pd.perturbation* function and calculated the change of total viral gene expression after perturbation.

#### Analyses of SARS-CoV-2 dataset

We subsampled the cells based on their infection status. We adapted the monocle preprocessing pipeline implemented in dynamo and excluded the UMI reads from lenti-virus. We obtained 5,001 infected cells and 1,000 highly variable genes for downstream analyses. We used both highly variable genes in host and virus genes to perform PCA and calculated the nearest neighbors with k = 30. Then 269 velocity genes were used for downstream analyses. We used the apoptosis-related genes collected by KEGG to calculate apoptosis activity score of each cell using *dynamo.tl.score_cells* function.

### Analyses of multi-modality datasets

#### RNA velocity estimation for multi-modality datasets

In traditional scRNA-seq datasets, RNA velocity methods use smoothed spliced and unspliced RNA counts through nearest-neighbor pooling, based on the PCA space computed from transcripts alone. However, this approach is not suitable for multimodal scenarios, as it overlooks hidden variables by relying on a single modality. To construct a consistent manifold combining information from multi-modal genomics data, we utilized WNN as implemented in MultiVelo^13^. The WNN algorithm combines low-dimensional representations from RNA and ATAC omics data. Specifically, we use PCA results from scRNA-seq data and latent semantic indexing (LSI) from scATAC-seq as inputs. The nearest neighbors identified by WNN were then used to calculate the first moment, reducing noise in separate modalities and approximating a unified manifold for GraphVelo.

#### Mouse skin dataset preprocessing

The preprocessed SHARE-seq mouse skin dataset^63^ is adopted directly from MultiVelo data resources. All the procedures are consistent with MultiVelo except we get the LSI representation processed by SCARlink^77^. We construct the WNN graph using 50 nearest neighbors for downstream calculation. We run scVelo with ‘stochastic’ mode to estimate the RNA velocity based on the WNN graph as we discussed above.

#### Human cortex dataset preprocessing

The preprocessed human cerebral cortex data is adopted directly from MultiVelo data resources. All the procedures are consistent with MultiVelo except we get the LSI representation processed by SCARlink. We construct the WNN graph using 50 nearest neighbors for downstream calculation. We run scVelo with ‘stochastic’ mode to estimate the RNA velocity based on the WNN graph.

### Data availability

All the sequencing raw data are publicly accessible. The A549 dataset can be accessed via https://figshare.com/ndownloader/files/53666738. The FUCCI cell cycle data can be downloaded from https://figshare.com/ndownloader/files/53705057. The mouse gastrulation subset to erythroid lineage can be extracted using scVelo’s CLI: *scvelo.datasets.gastrulation_erythroid()* or from the original work under accession number E-MTAB-6967 of ArrayExpress. The human bone marrow can be extracted using scVelo’s CLI: *scvelo.datasets.bonemarrow()* or through the Human Cell Atlas data portal at https://data.humancellatlas.org/explore/projects/091cf39b-01bc-42e5-9437-f419a66c8a45. The mouse coronal hemibrain spatial transcriptomic data can be downloaded from (https://www.dropbox.com/s/c5tu4drxda01m0u/mousebrain_bin60.h5ad?dl=0). The original HCMV infected moDC data can be accessed via Zenodo (https://zenodo.org/records/10404879) and the processed data can be downloaded from https://figshare.com/ndownloader/files/53666756. The SARS-CoV-2 data can be accessed via https://figshare.com/ndownloader/files/53666588. The preprocessed mouse skin development dataset can be accessed via https://figshare.com/articles/dataset/Mouse_Hair_Follicle_RNA_Data/22575307 and https://figshare.com/articles/dataset/Mouse_hair_follicle_ATAC_data/22575313. The preprocessed human cortex dataset can be downloaded from https://figshare.com/articles/dataset/Developing_Human_Cortex_RNA_Data/22575376 and https://figshare.com/articles/dataset/Developing_Human_Cortex_ATAC_Data/22575370.

### Code availability

Python package GraphVelo can be accessed from https://github.com/xing-lab-pitt/GraphVelo. Reproducibility and tutorials can be found in https://github.com/xing-lab-pitt/GraphVelo/tree/main/notebook and https://graphvelo.readthedocs.io/en/latest/.

## Author contributions

JX conceived and formulated the theoretical framework, YC, and JG performed tests on simulated data, YC and JX analyzed scRNA-seq datasets, YC, YZ, JG, and KN developed the package GraphVelo, YC and JX wrote the draft, and all authors edited the manuscript.

## Acknowledgments

This work was partially supported by National Institute of General Medical Sciences (1R01GM148525 to JX and 1R01GM139297 to IB), and National Science Foundation (2325149) to JX.

This work was supported by the National Key Research and Development Program of China (2023YFE0112300); National Natural Sciences Foundation of China (32261133526; 32270709; 32070677); the 151 talent project of Zhejiang Province (first level), the Science and Technology Innovation Leading Scientist (2022R52035)

## Figures

**Extended Data Fig. 1.**
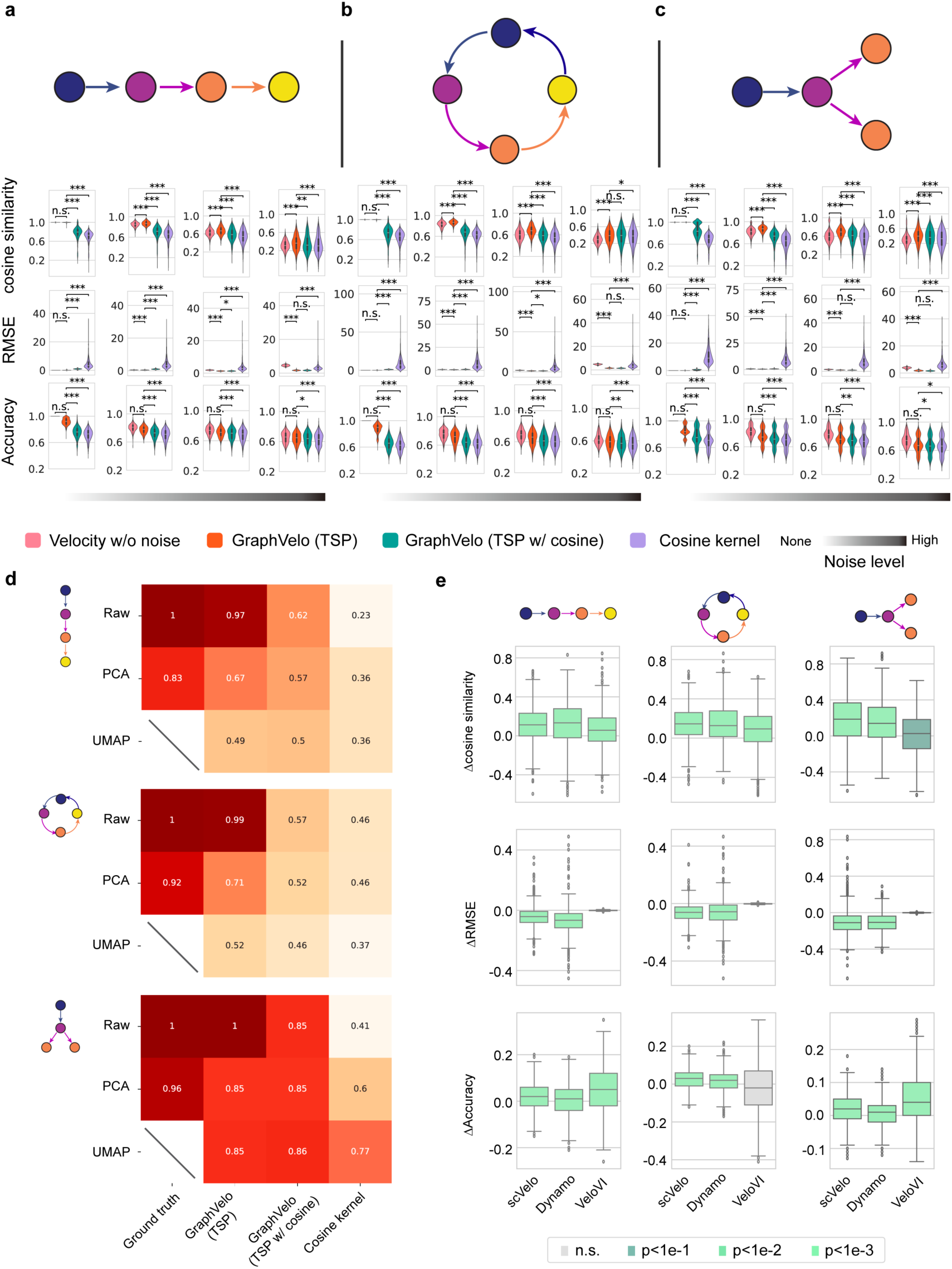
Refining RNA velocity from noisy simulation data or tradition splicing-based methods. **(a-c)** Evaluation of on simulated scRNA-seq data under linear, cycling and bifurcating differentiation models with an increasing noise level in velocity vectors. **(d)** Heatmap of the correlation of cell speed calculated as the norm of velocity vector with respect to the full-dimensional RNA velocity and the norm of velocity vectors projected to PCA or UMAP space using GraphVelo, GraphVelo with cosine regularization, cosine kernel. **(e)** Boxplots of metric evaluations on GraphVelo correction to the original velocities estimated by scVelo, dynamo, and VeloVI. Box colors indicate the statistical test results for improvement relative to zero.

**Extended Data Fig. 2.**
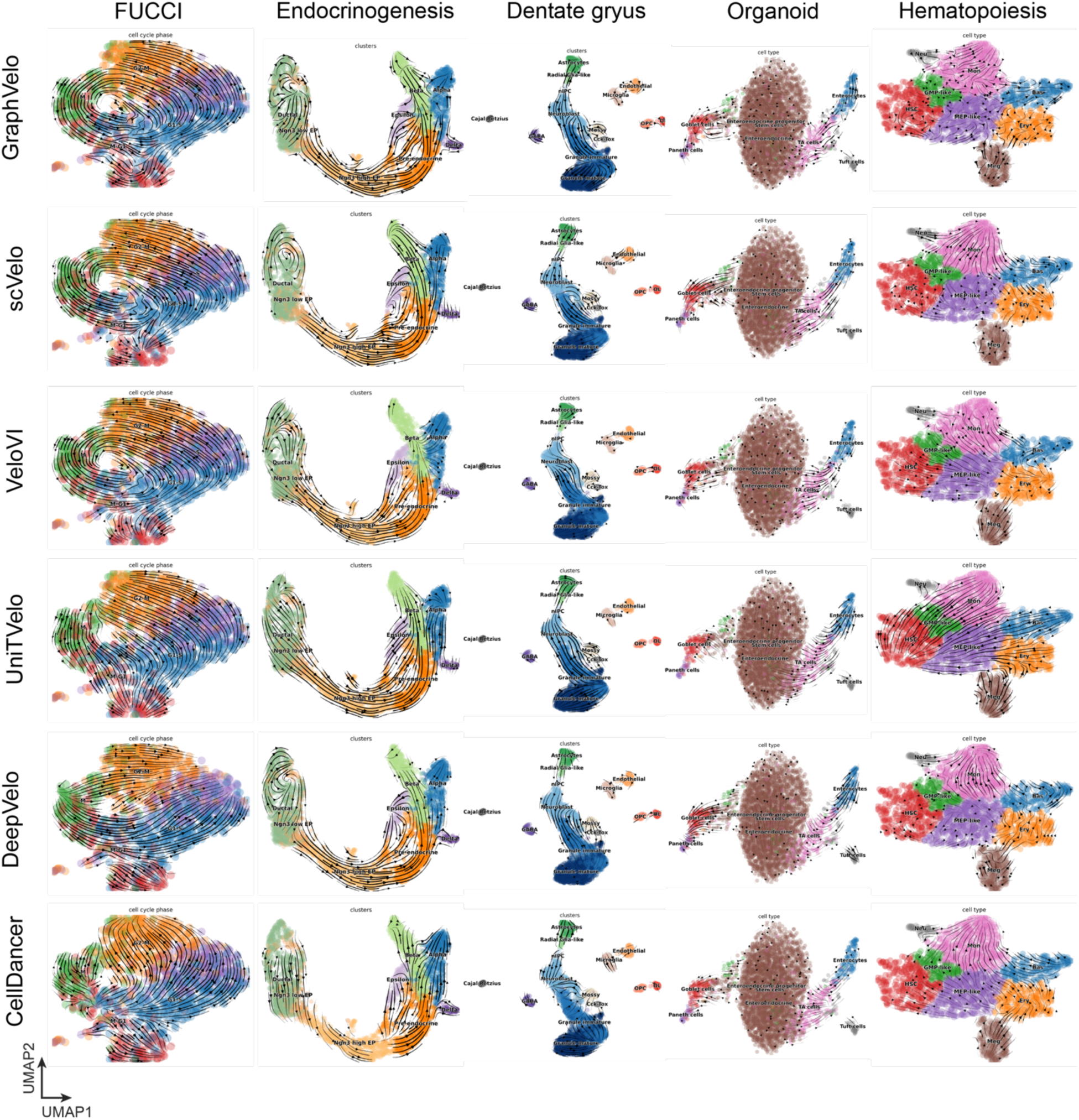
The visualization results of inferred cell fate transition directions in UMAP spaces.

**Extended Data Fig. 3.**
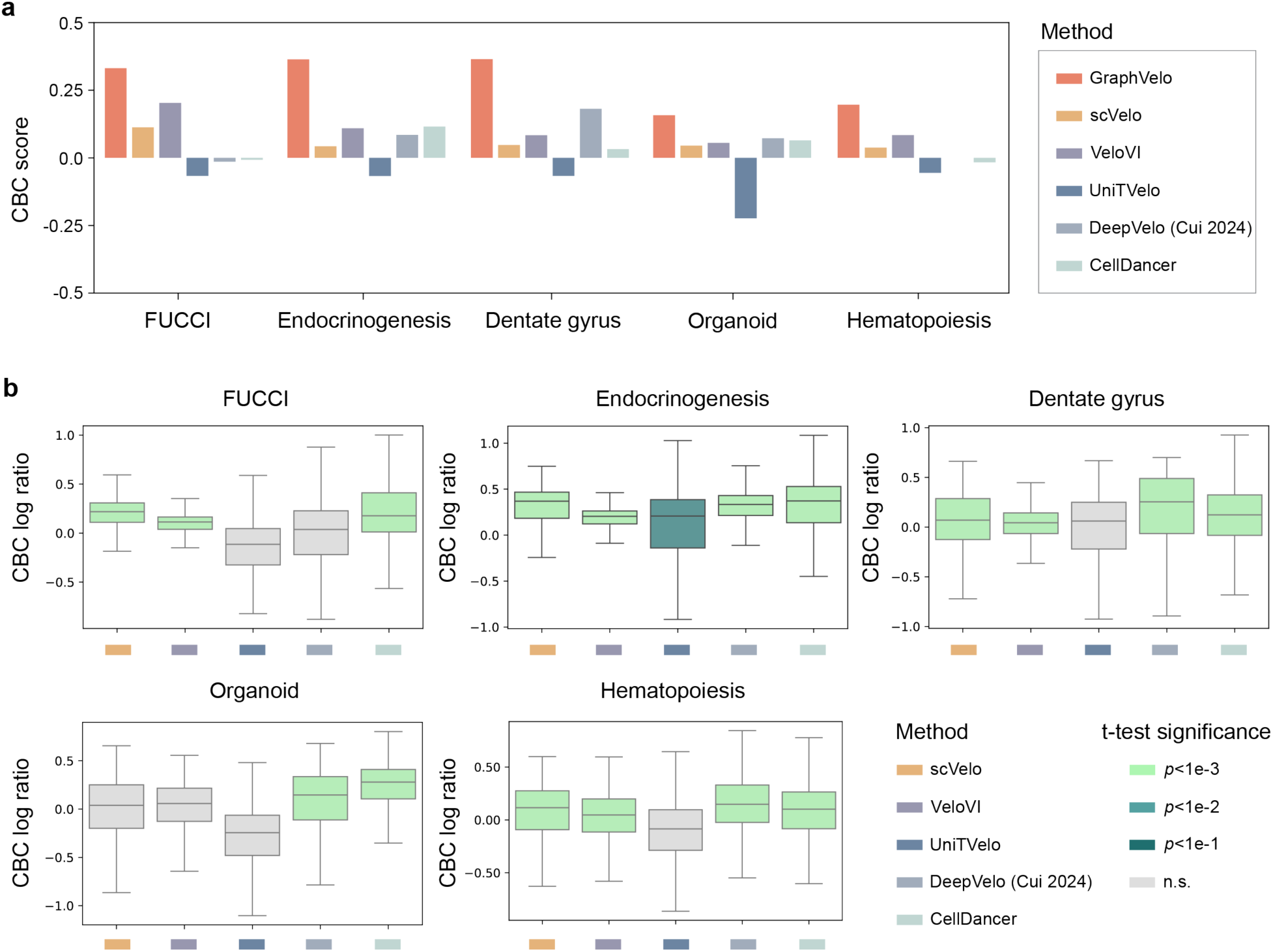
Quantitative benchmarks on GraphVelo across diverse biological systems. **(a)** The average CBC scores across five benchmark datasets using different RNA velocity estimation methods. **(b)** Pairwise comparisons before and after applying GraphVelo using the outputs of distinct RNA velocity algorithms across datasets. The Welch’s t-test is performed to demonstrate a significant improvement of applying GraphVelo against zero.

**Extended Data Fig. 4.**
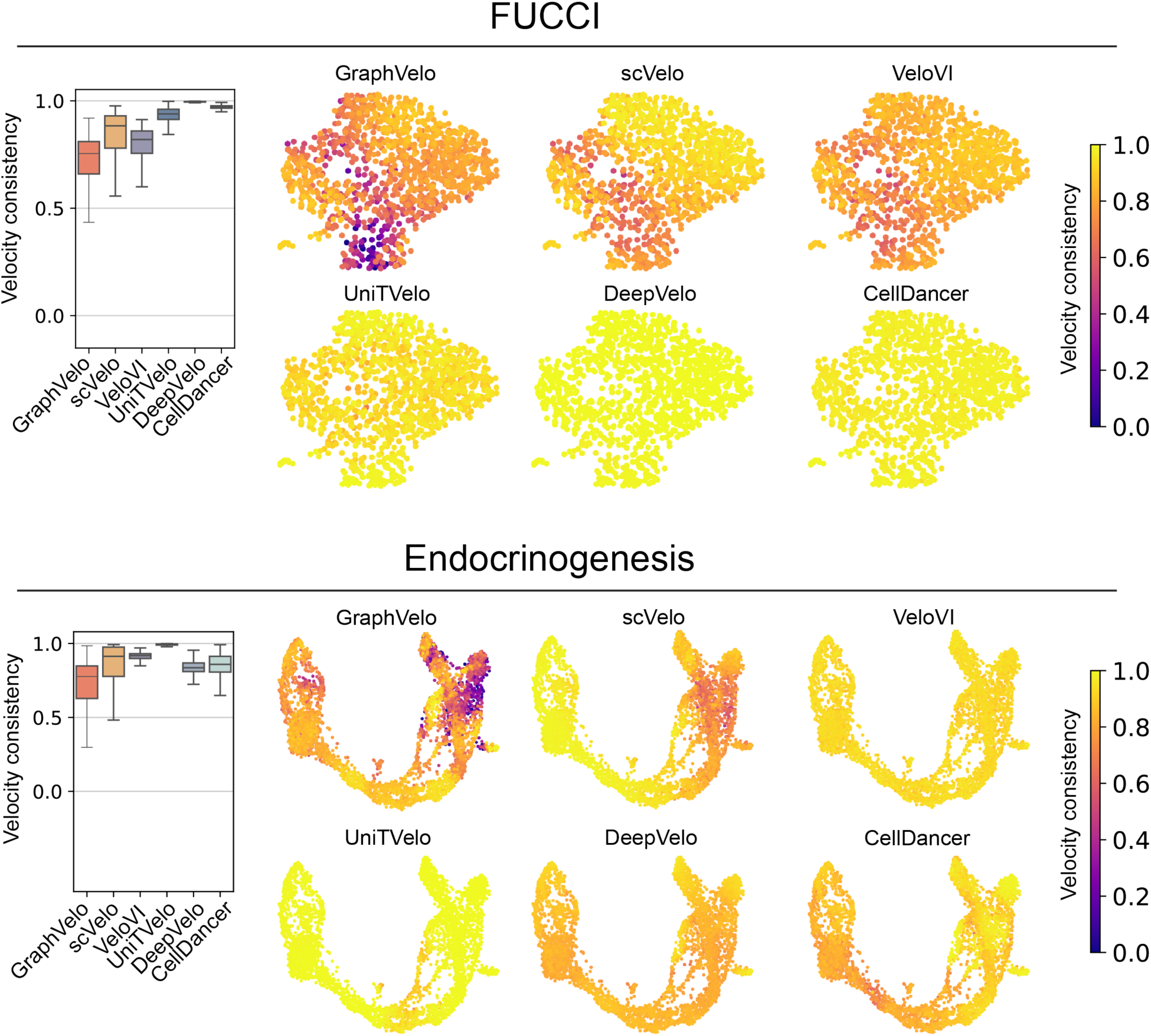
Benchmarking velocity consistency on FUCCI and endocrinogenesis dataset.

**Extended Data Fig. 5.**
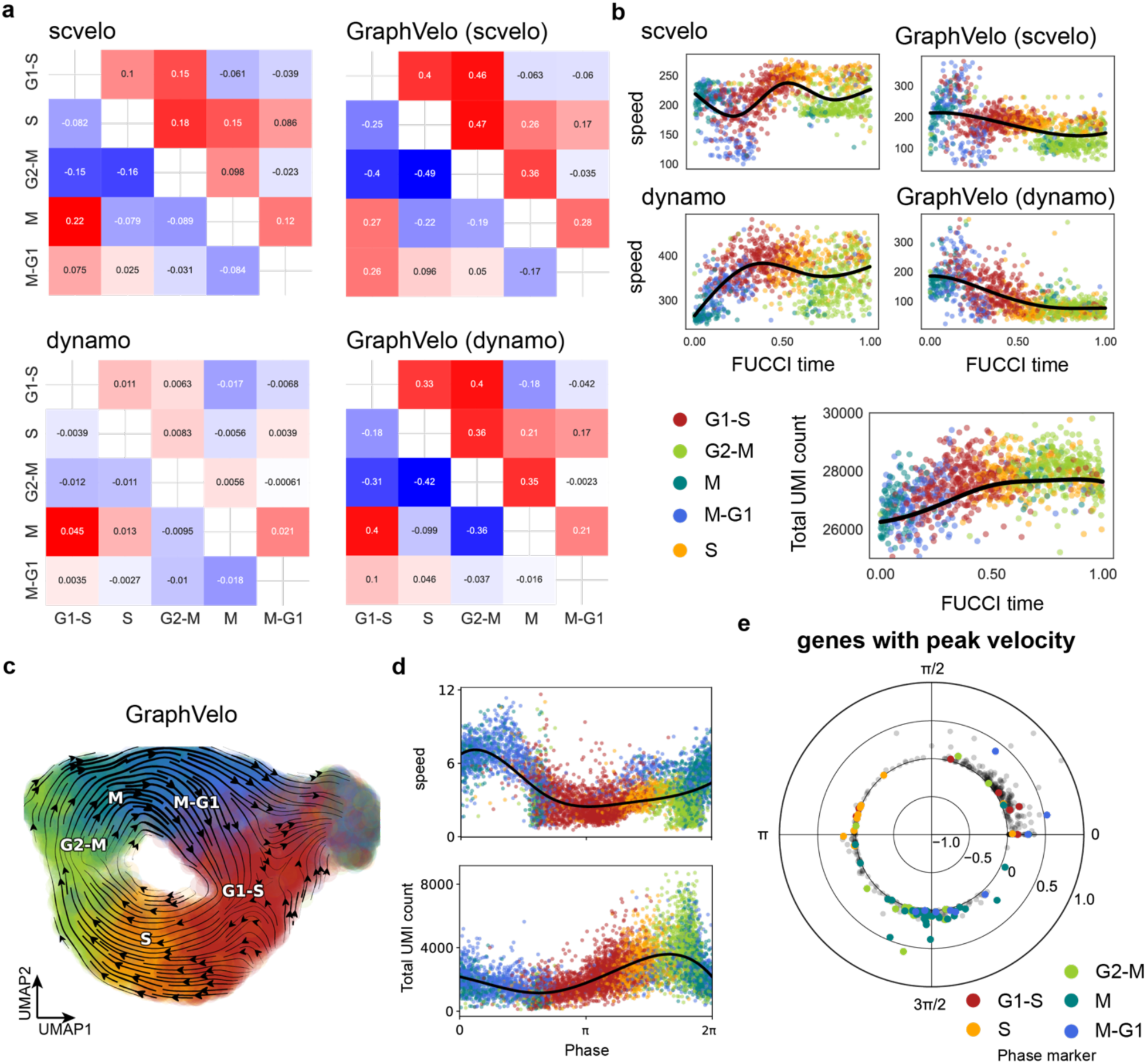
Quantitative evaluation of GraphVelo on two independent cell cycle datasets. **(a)** Pairwise CBC scores between cell cycle phases (G1-S, S, G2-M, M, M-G1) using velocity vectors estimated by scVelo, dynamo, and their GraphVelo-refined outcomes. **(b)** Cell cycle speed comparison along FUCCI time. GraphVelo-refined velocities identify the peak region within M-G1 phase and show a better match to total UMI distribution, reflecting more accurate scaling of cell cycle dynamics. **(c)** Vector field reconstructed by GraphVelo on the A549 dataset, projected onto UMAP. Cells are colored by their cell cycle phases. **(d)** Cell cycle speed (top) and total UMI count changes (bottom) plotted along the cell cycle phase. Peaks in velocity magnitude coincide with result in FUCCI data, particularly in M-G1. **(e)** Polar plot of genes with peak RNA velocity, stratified by cell cycle phase, reflecting the cascade activation of phase marker genes.

**Extended Data Fig. 6.**
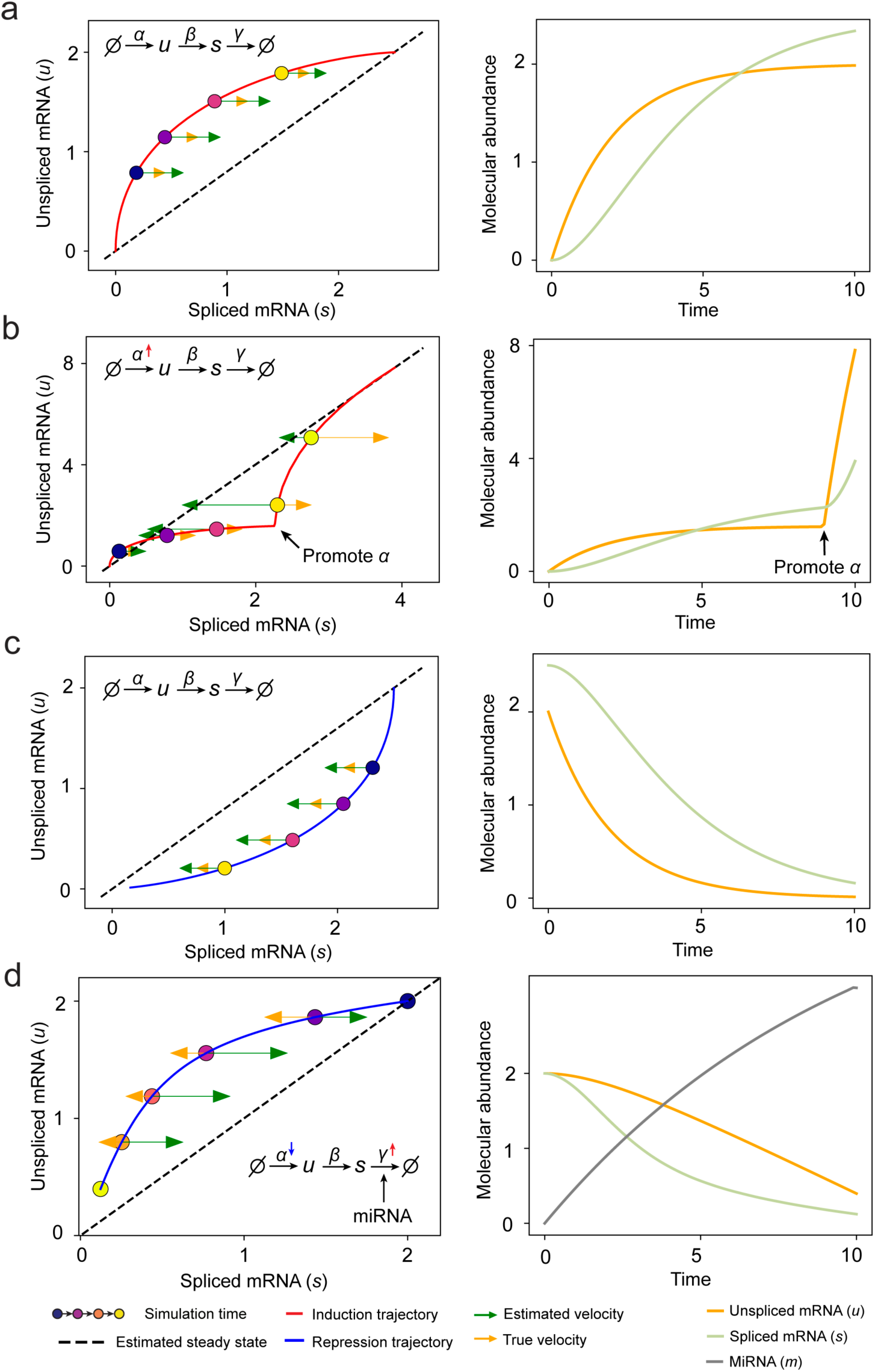
RNA velocity inference on simulation results of splicing kinetics with constant or changed rates. **(a)** Standard splicing kinetics along an induction trajectory in phase portrait and the reactions define how abundance levels of molecules change along simulation time. Right panel shows corresponding trajectories over time (same below). **(b)** Transcription burst along an induction trajectory in phase portrait. The transcription rate constant ***α*** was promoted at specific time point. **(c)** Standard splicing kinetics along a repression trajectory. **(d)** Rapid time-varying degradation kinetics along repression trajectory. External signal promotes synthesis of microRNA, which enhances degradation of target mRNA resulting in a microRNA - dependent varying degradation rate “constant” ***γ***, and inhibits the target gene via a decreasing ***α***.

**Extended Data Fig. 7.**
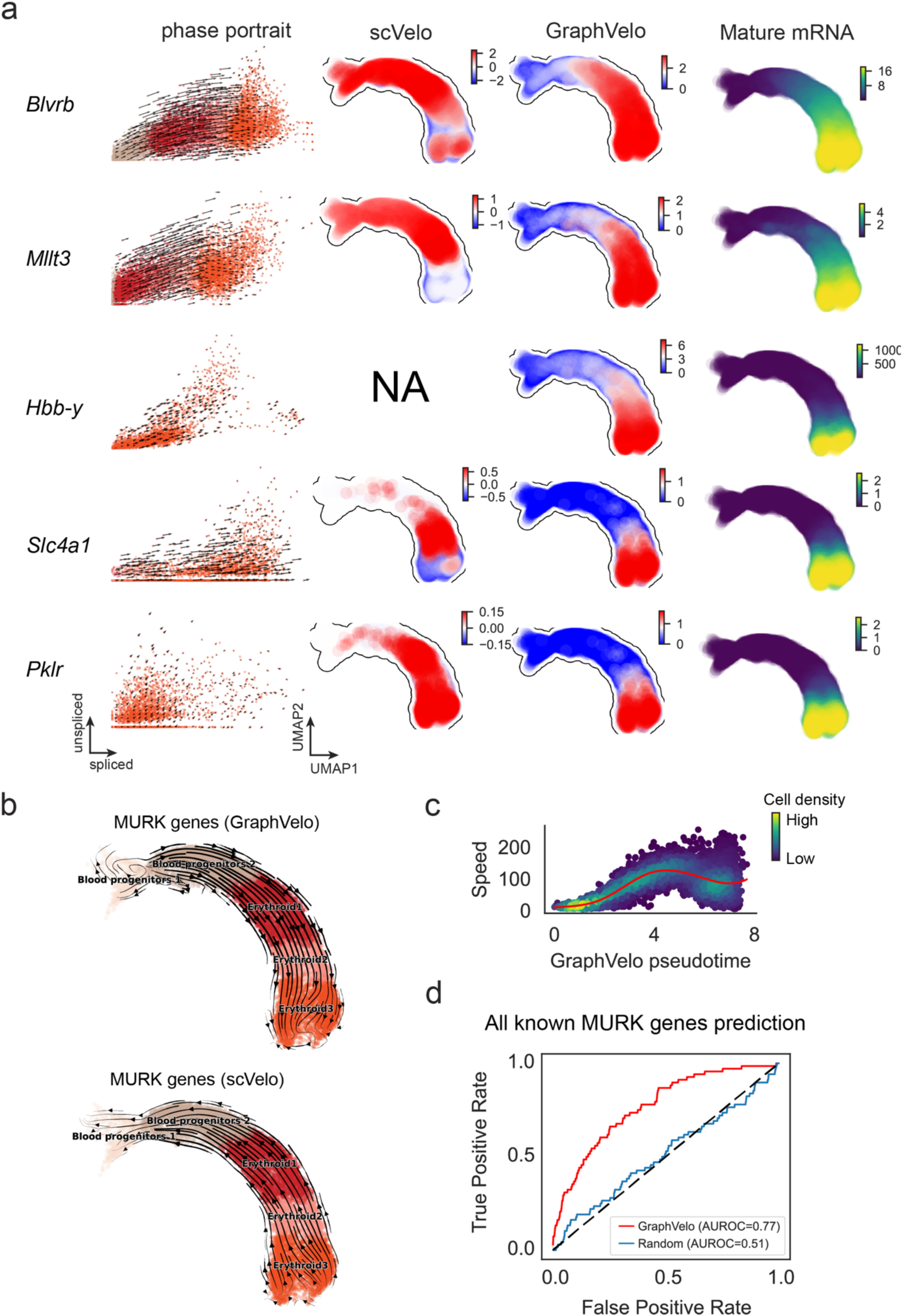
GraphVelo extrapolation of RNA velocities of an extended list of gene set in mouse erythroid maturation dataset. **(a)** Phase portrait with GraphVelo vectors, velocity estimated by scVelo, refined velocity by GraphVelo, and gene expression of mature mRNA of a selected set of genes. Cells were colored by cell type, corresponding velocity, and mature mRNA abundance, respectively, and visualized on the phase portrait and UMAP, respectively. **(b)** Velocities of MURK genes derived from GraphVelo and scVelo for gastrulation erythroid maturation cells projected to a predefined UMAP representation. Note that RNA velocity estimated by scVelo was in a reverse flow, possibly influenced by transcription burst events. **(c)** Cell speed distribution along the vector field pseudotime axis. Cells were colored by local density and red line indicates the fitted curve. **(d)** Receiver operating curve analyses of MURK gene prediction based on dynamo accelerations analysis using GraphVelo as input, in contrast to a random predictor. AUC, area under curve.

**Extended Data Fig. 8.**
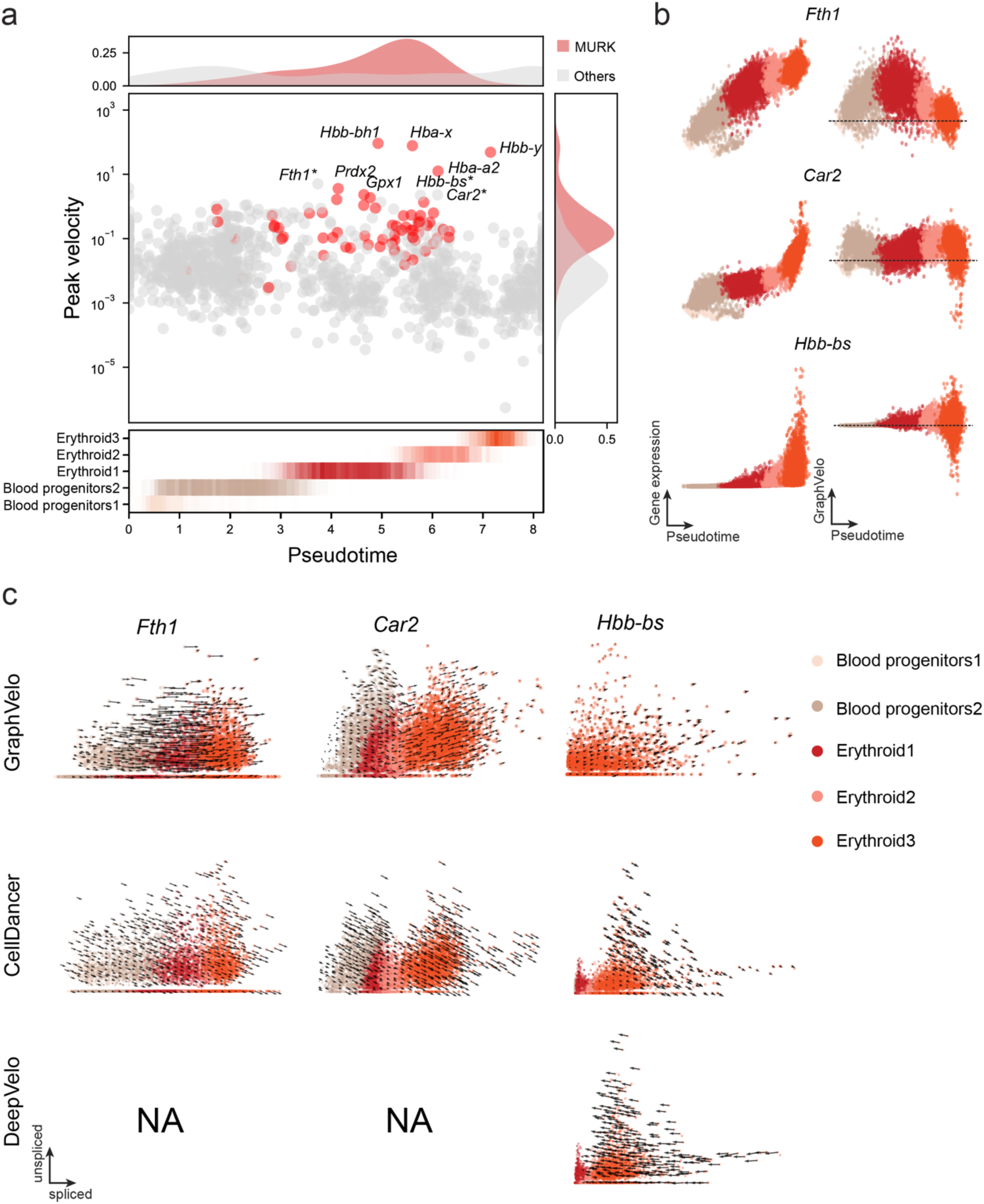
GraphVelo reveals potential MURK genes with larger peak velocity magnitude during erythroid maturation. **(a)** Peak RNA velocity across pseudotime for highly variable genes. MURK genes (red) show significantly higher peak velocity magnitudes than other genes (gray), particularly in late-stage erythroid cells. Top peak velocity genes are annotated and *Fth1*, *Car2*, *Hbb-bs* are suspected as potential MURK genes and highlighted by star*. Density plots above and to the side illustrate the distributions of pseudotime and peak velocity, respectively. The heatmap below shows cell density along pseudotime by cell states. **(b)** Expression and corresponding velocities of *Fth1*, *Car2*, *Hbb-bs* along the vector field-based pseudotime in mouse erythroid data. **(c)** Phase portraits comparing velocity vectors estimated by GraphVelo, CellDancer, and DeepVelo for selected potential MURK genes. GraphVelo reveals well-directed and temporally aligned transcriptional dynamics, while alternative methods (particularly DeepVelo) fail to provide interpretable velocity fields for key genes with altered kinetics.

**Extended Data Fig. 9.**
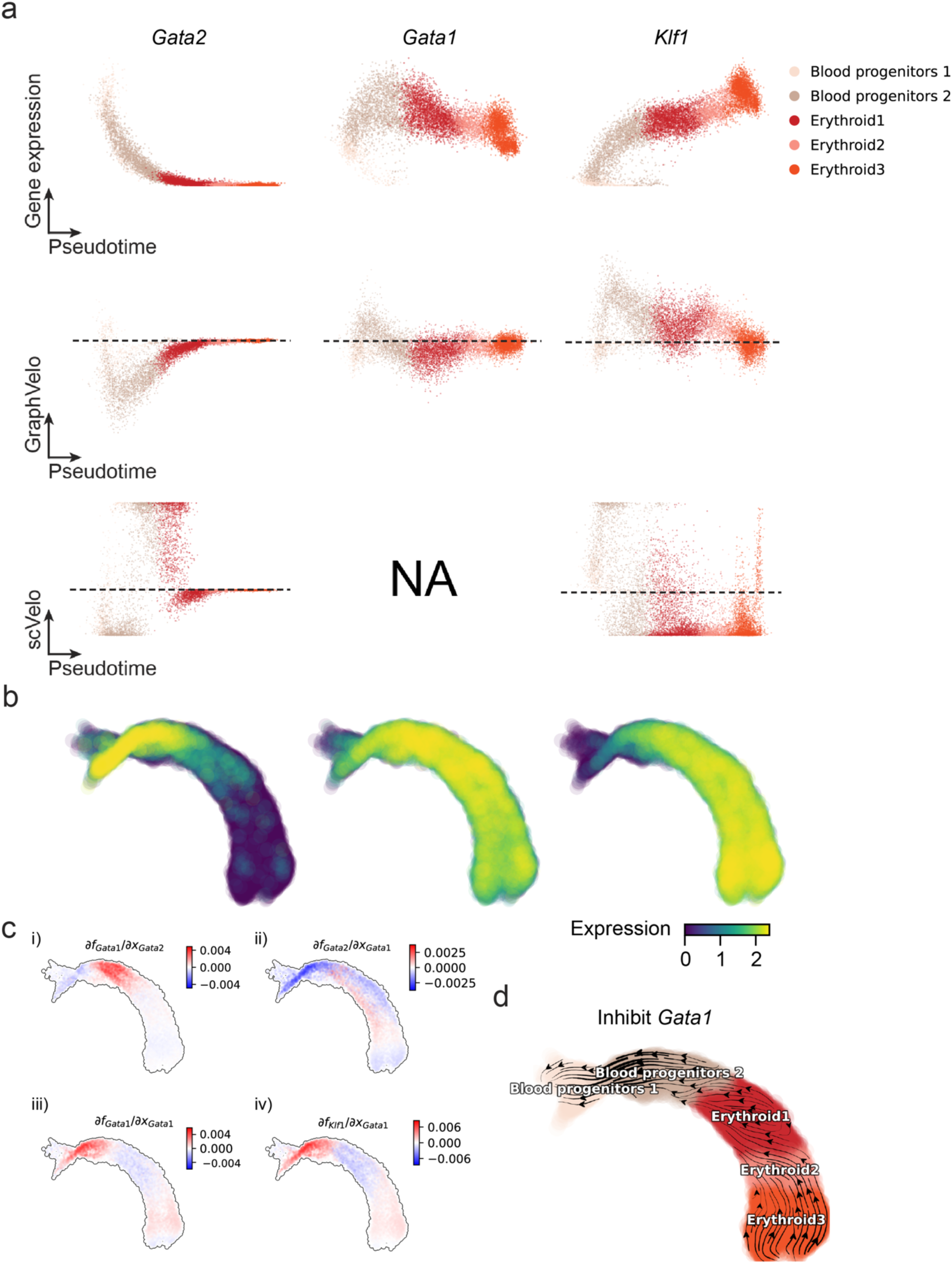
Regulatory cascades of driver TFs in mouse erythroid data. **(a)** Expression and corresponding velocities of *Gata2*, *Gata1*, *Klf1* along the vector field-based pseudotime in mouse erythroid data. **(b)** Gene expression of *Gata2*, *Gata1*, *Klf1* on the UMAP space. **(c)** Molecular mechanisms of driver TFs underlying erythroid lineage commitment based on Jacobian analyses. (i) *Gata2* activates *Gata1*. (ii) Repression of *Gata2* by *Gata1*. (iii) Self-activation of *Gata1*. (iv) *Gata1* activates *Klf1*. **(d)** In silico perturbation analyses on GraphVelo-based vector field to examine the role of *Gata1* in gastrulation erythroid maturation.

**Extended Data Fig. 10.**
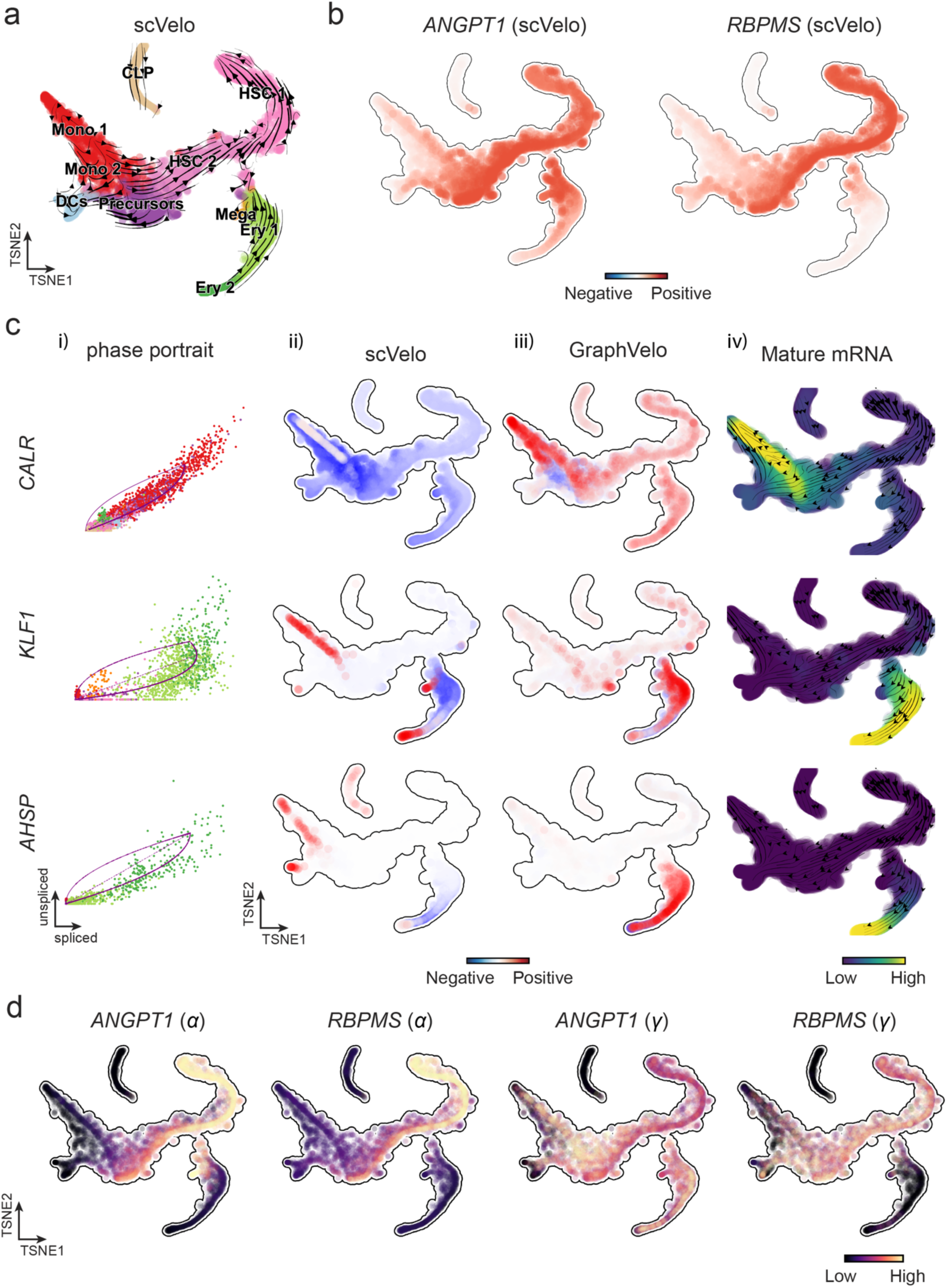
RNA velocity estimated by scVelo and refined by GraphVelo from a hematopoiesis dataset. **(a)** Velocities derived from scVelo in the hematopoiesis development and projected to a pre-defined TSNE embedding. **(b)** TSNE visualization of the RNA velocity estimated by scVelo of *ANGPT1* and *RBPMS* genes. **(c)** Scatter plots of: i) phase portrait, ii) velocities estimated by scVelo, iii) refined velocities by GraphVelo, and iv) mature mRNA expression of transcription burst genes (e.g. *CALR*, *KLF1* and *AHSP*). Cells were colored by cell type, corresponding velocity, and mature mRNA abundance, respectively, and visualized on the phase portrait and TSNE, respectively. **(d)** Cell-specific transcription rate constant ***α*** and degradation rate constant ***γ*** of rapid degradation genes *ANGPT1* and *RBPMS* visualized on the TSNE space, which is consistent with simulation result in Extended Data Fig. 3d.

**Extended Data Fig. 11.**
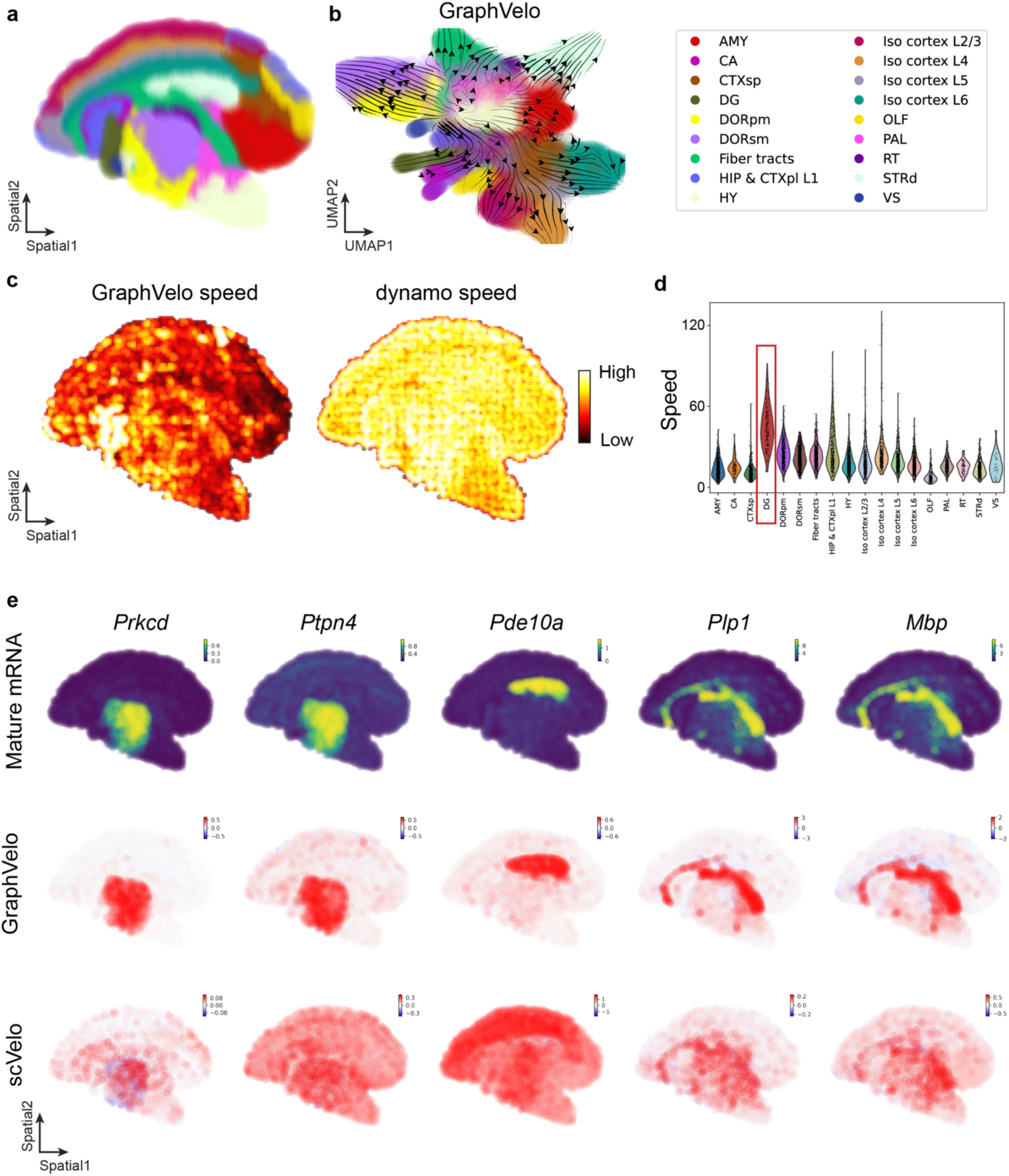
GraphVelo infers quantitative transcription rate from mouse coronal hemibrain spatial data. **(a)** The spatial location of cells colored with annotation in the original research. **(b)** UMAP visualization of the RNA velocity estimated by GraphVelo. **(c)** The transcription speed inferred by GraphVelo (left panel) and dynamo (right panel). **(d)** Violin plot of cell-wise transcription speed grouped by cell state. **(e)** Spatial distribution of mature mRNA expression, velocities estimated by dynamo, and refined velocities by GraphVelo of spatial variable genes. Cells were colored by mature mRNA abundance and corresponding velocity, respectively, and visualized on spatial location.

**Extended Data Fig. 12.**
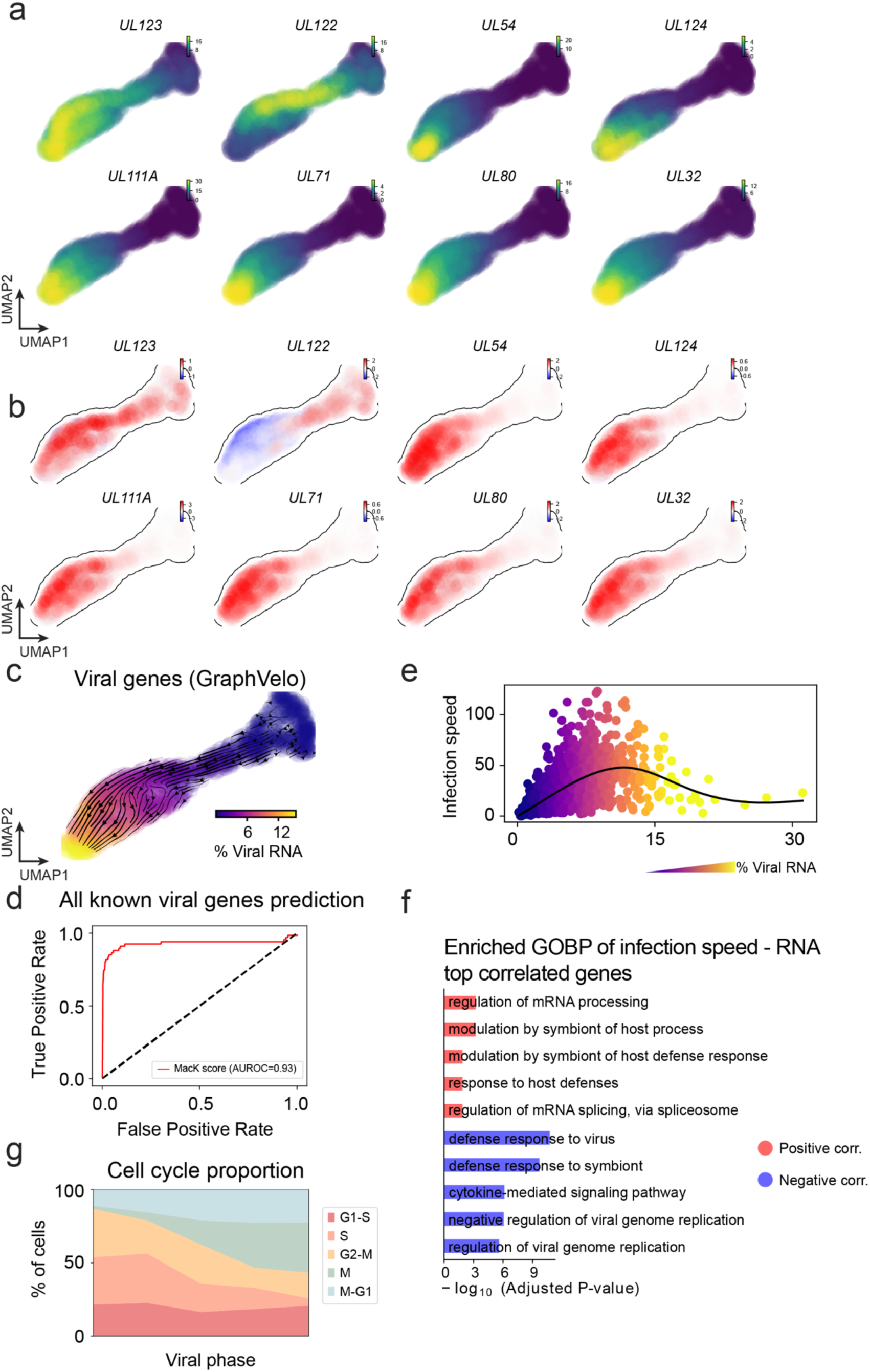
Dynamic patterns of gene trajectories during HCMV infection. **(a)** Gene expression of selected viral factors on the UMAP space. **(b)** Corresponding velocities inferred by GraphVelo for viral factors in (a). **(c)** Viral RNA velocities derived from GraphVelo for infected cells, projected to the UMAP representation only using viral genes. **(d)** Receiver operating curve (ROC) analyses of MacK scores when using all detected viral genes as the gold standard. **(e)** Viral infection speed versus viral load. Cells were colored by the percentage of viral RNA. **(f)** GO enrichment analyses of top host genes correlated with infection speed. **(g)** Shifts in cell proportions across different cell cycle phases along the viral phase trajectory. Cells were divided into bins according to viral RNA percentage. The result indicates a decreasing S phase population corresponding to infection-driven transcriptional alterations of the cell cycle.

**Extended Data Fig. 13.**
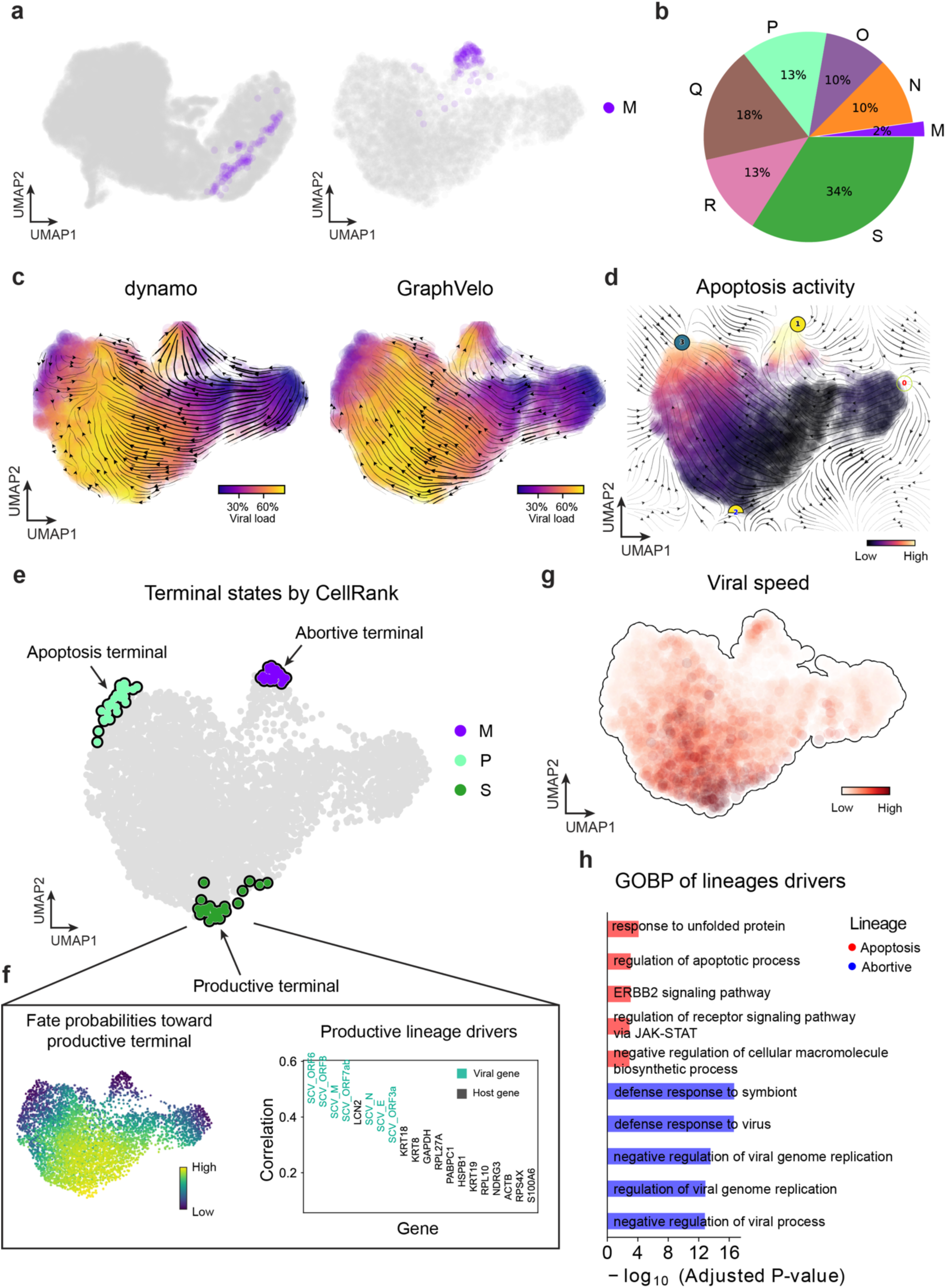
GraphVelo infers multiple terminals driven by viral-host interactions. **(a)** UMAP coordinates computed from all cell populations in the original study^55^ (left) and from the infected subpopulation alone (right). Cells belonging to cluster M are highlighted in purple. **(b)** Proportion of cluster M within all infected cell population. **(c)** Velocities derived from dynamo (left) and GraphVelo (right) in the infected cells and projected to UMAP embedding. Cells are colored by viral load. **(d)** Topological analyses of GraphVelo vector field identified initial position with low viral RNA load, saddle point with high viral load (as shown in panel c) and attractors residing in terminal states with high apoptosis activity. **(e)** Terminal states identified by CellRank using GraphVelo-corrected RNA velocity. **(f)** Fate probabilities toward the productive terminal state (left) and top 20 lineage-correlated genes identified (right) based on GraphVelo-based CellRank analysis. **(g)** GO enrichment analyses of top lineage-driver genes along two distinct cell death trajectories.

**Extended Data Fig. 14.**
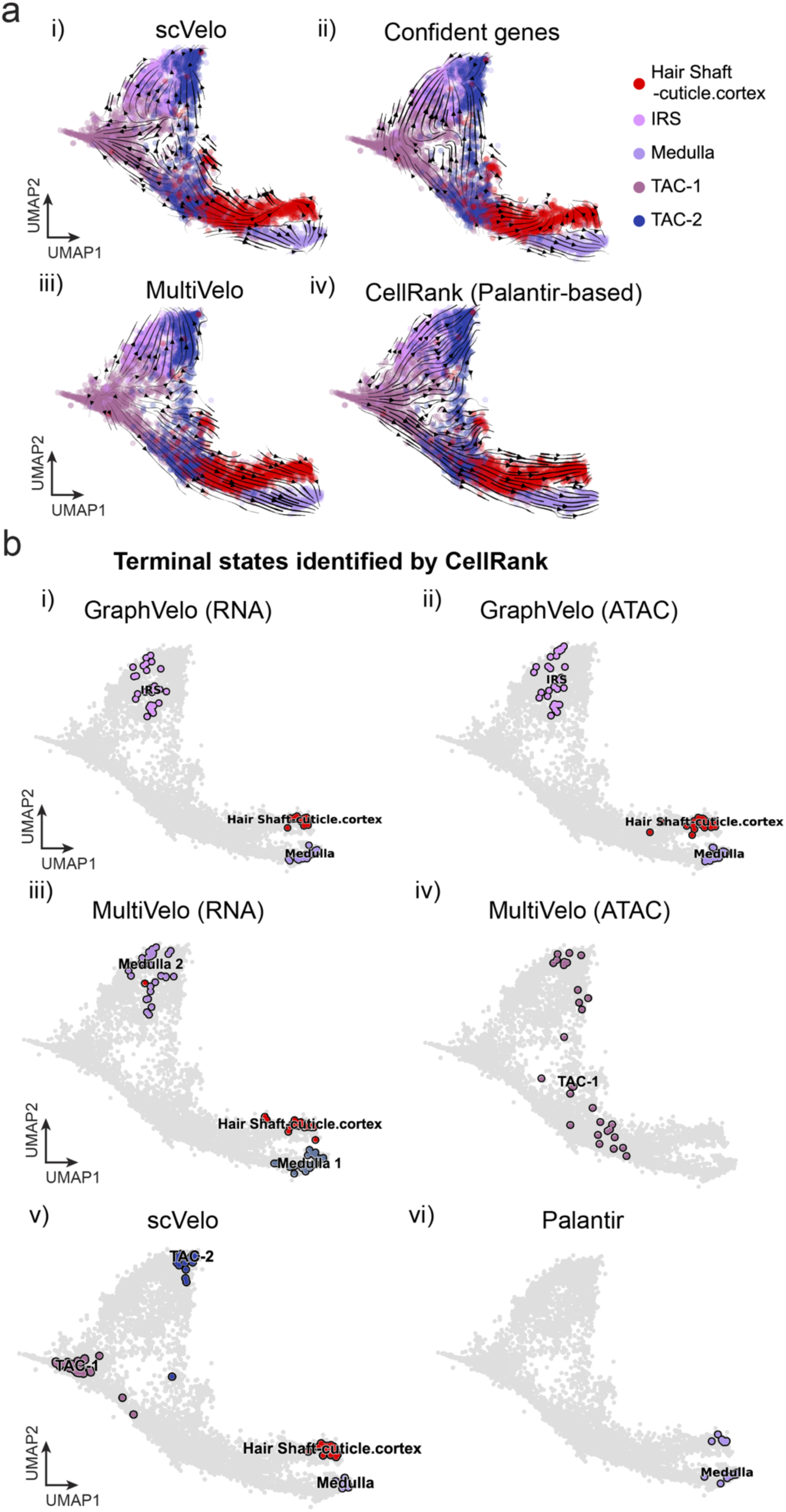
GraphVelo estimation of modality-consistent vector field based on multi-omics velocities. **(a)** Velocity field projected to the pre-defined UMAP representation with different methods or gene sets: i) scVelo RNA velocity based on all velocity genes; ii) scVelo RNA velocity based on confident velocity genes filterred by dynamo criteria; iii) MultiVelo RNA velocity based on all velocity genes; iv) CellRank pseudotime kernel based on Palantir pseudotime. **(b)** Terminal states identified by CellRank using different representation and corresponding velocity: i) GraphVelo-corrected RNA velocity; ii) GraphVelo-computed chromatin velocity; iii) MultiVelo-computed RNA velocity; iv) MultiVelo-computed chromatin velocity; v) scVelo-computed RNA velocity; vi) Palantir-based pseudotime kernel.

**Extended Data Fig. 15.**
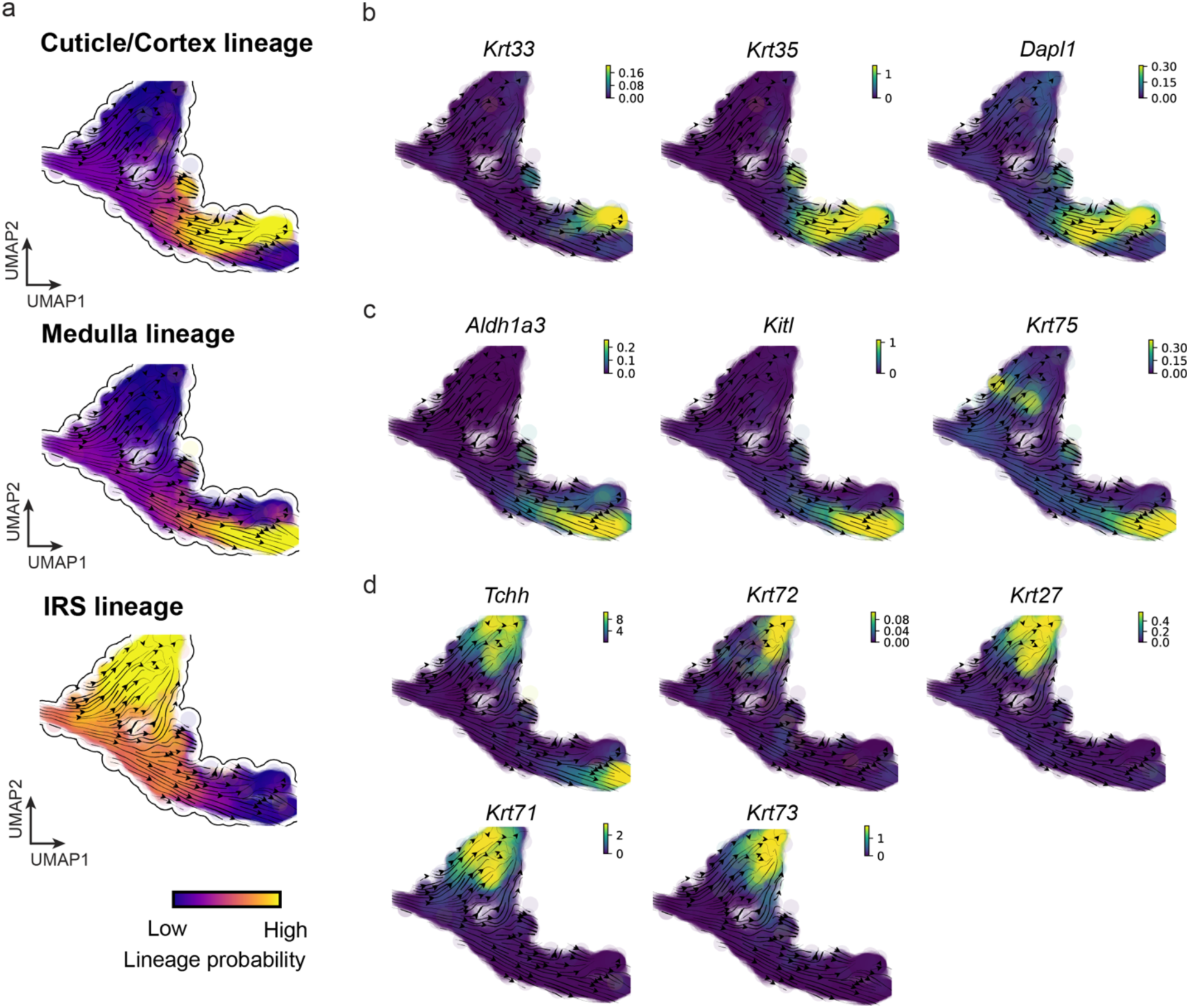
Reconstructed lineage commitment during mouse hair follicle differentiation using GraphVelo velocities. **(a)** Lineage commitment probablity of each terminal cell type on the UMAP vector field. **(b-d)** Gene expression distribution of markers for cuticle/cortex, medulla and IRS lineages on the UMAP vector field.

**Extended Data Fig. 16.**
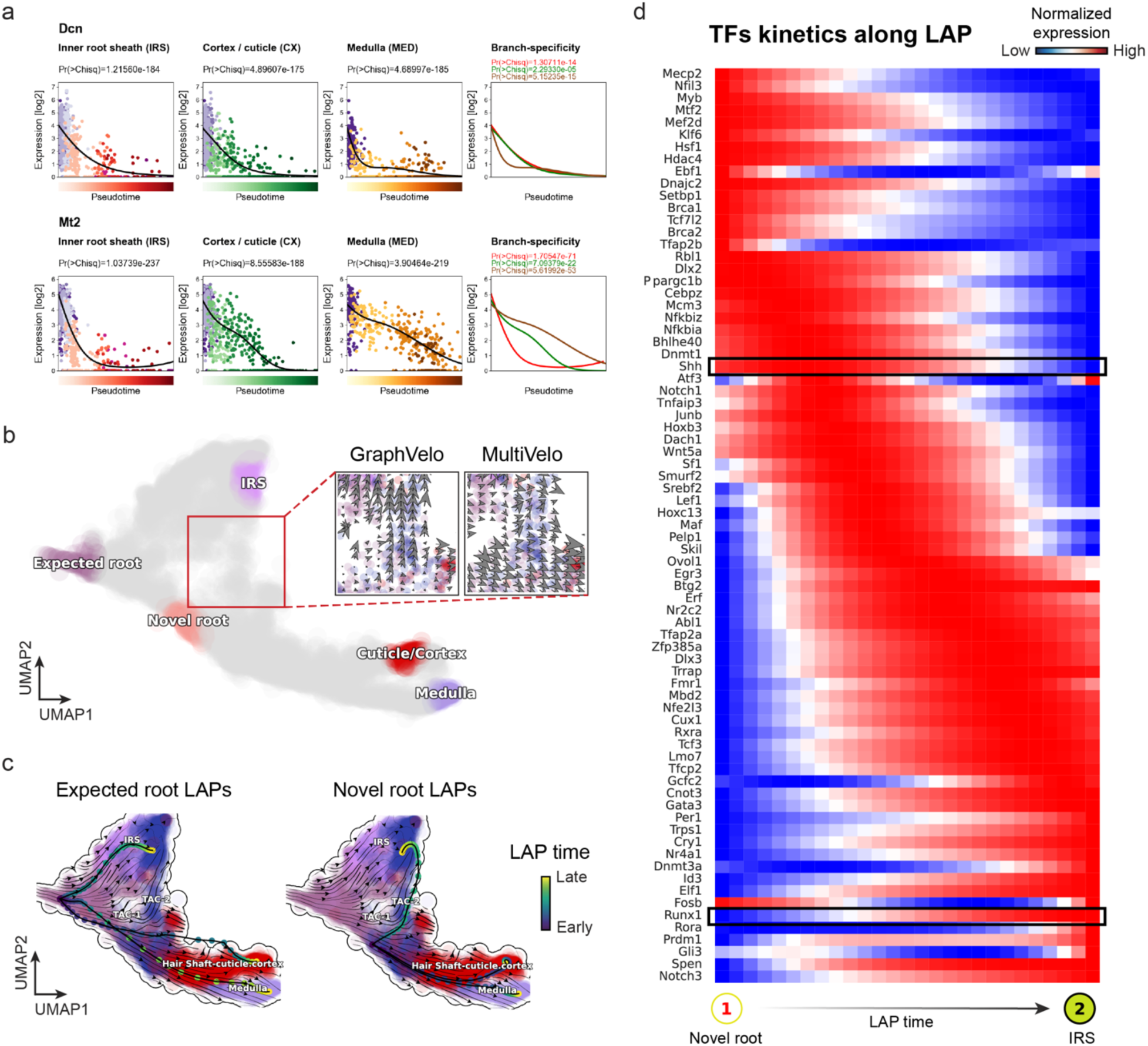
Novel root identified with multi-omic vector field from GraphVelo velocities. **(a)** *Dcn*, *Mt2* expression dynamics during anagen hair follicle keratinocytes^62^. **(b)** Regions identified by topological analysis. Insert are ummarized cell-state transition vectors calculated by GraphVelo and MultiVelo along the path from the novel root to IRS and projected onto the UMAP representation. **(c)** Predicted developmental LAPs from expected root or novel root to to each of the terminal cell types in the UMAP embedding. Color of the node along the paths indicates the LAP transition time. **(d)** TF expression profiles along the LAP from novel root to IRS. *Shh* decays, alongside the induction of the *Runx1* gene.

**Extended Data Fig. 17.**
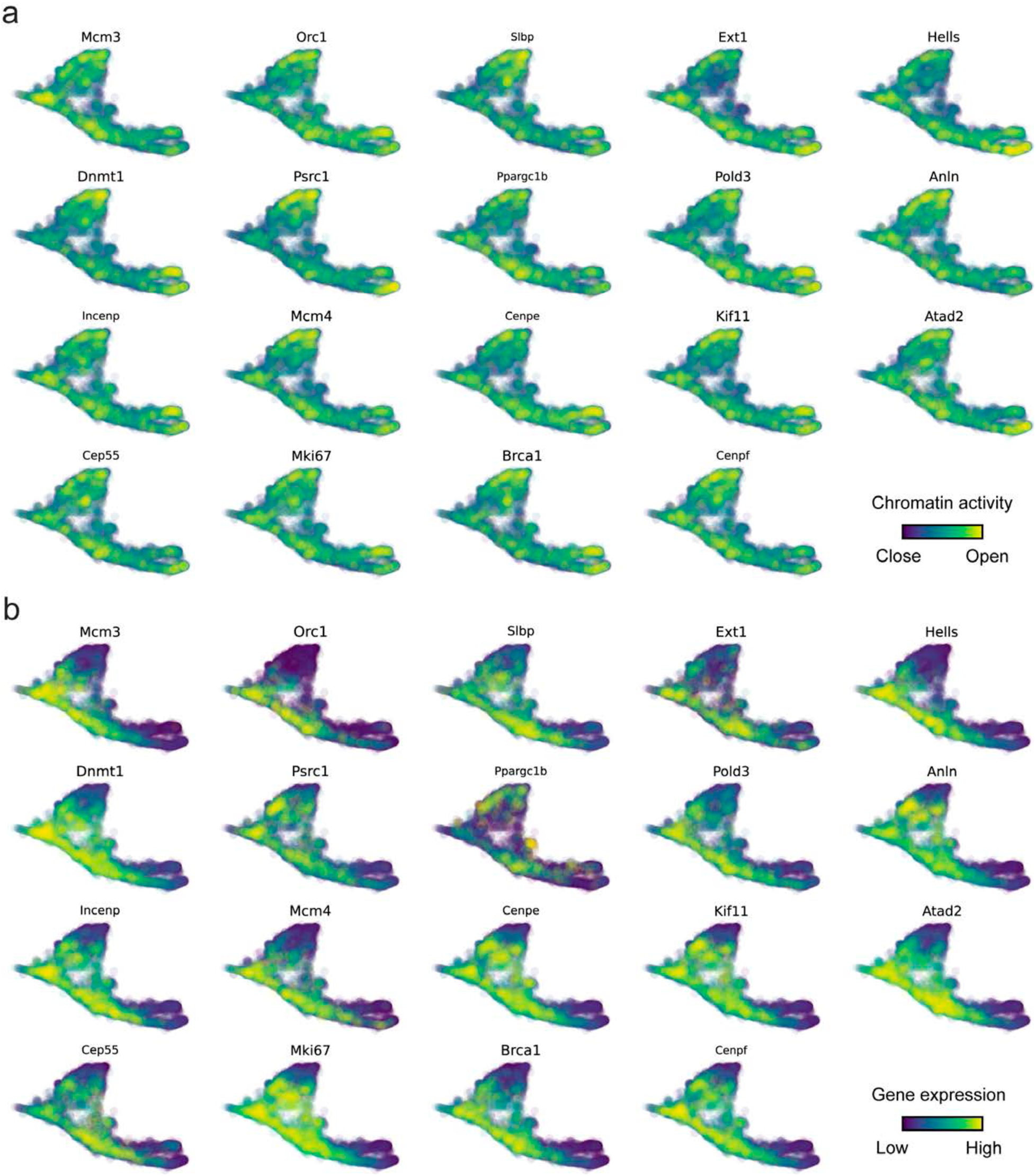
Genomic patterns of decoupled CCD genes. **(a)** Chromatin activity of decoupled CCD genes. **(b)** Gene expression of decoupled CCD genes.

**Extended Data Fig. 18.**
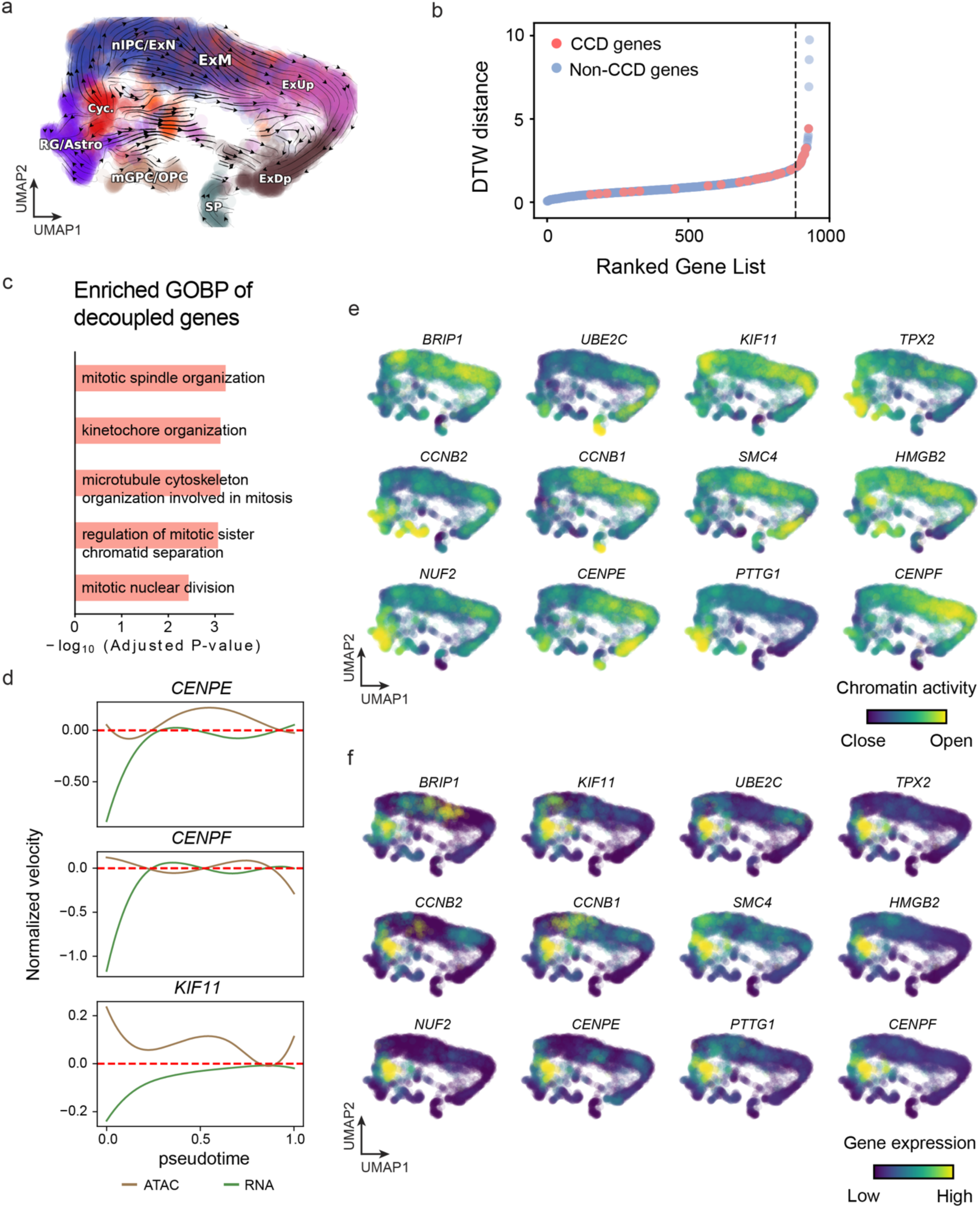
GraphVelo inferrence on epigenome and transcriptome decoupling dynamics in fetal human brain. **(a)** GraphVelo velocity field colored by cell macrostates. **(b)** DTW distance between RNA velocity and chromatin velocity of individual genes as a measure of the coupling/decoupling status. CCD genes were colored in red. The dotted line indicates the elbow point, with the decoupled genes on its right. **(c)** GO enrichment of decoupled genes in (c). **(d)** Line plot of nomarlized RNA and chromatin velocity along dynamo vector field pseudotime for genes predicted by GraphVelo to have notable decoupling patterns. Chromatin velocity trends were colored in brown and RNA velocity trends were colored in green. **(e)** Chromatin activity of decoupled CCD genes. **(f)** Gene expression of decoupled CCD genes.

**Extended Data Fig. 19.**
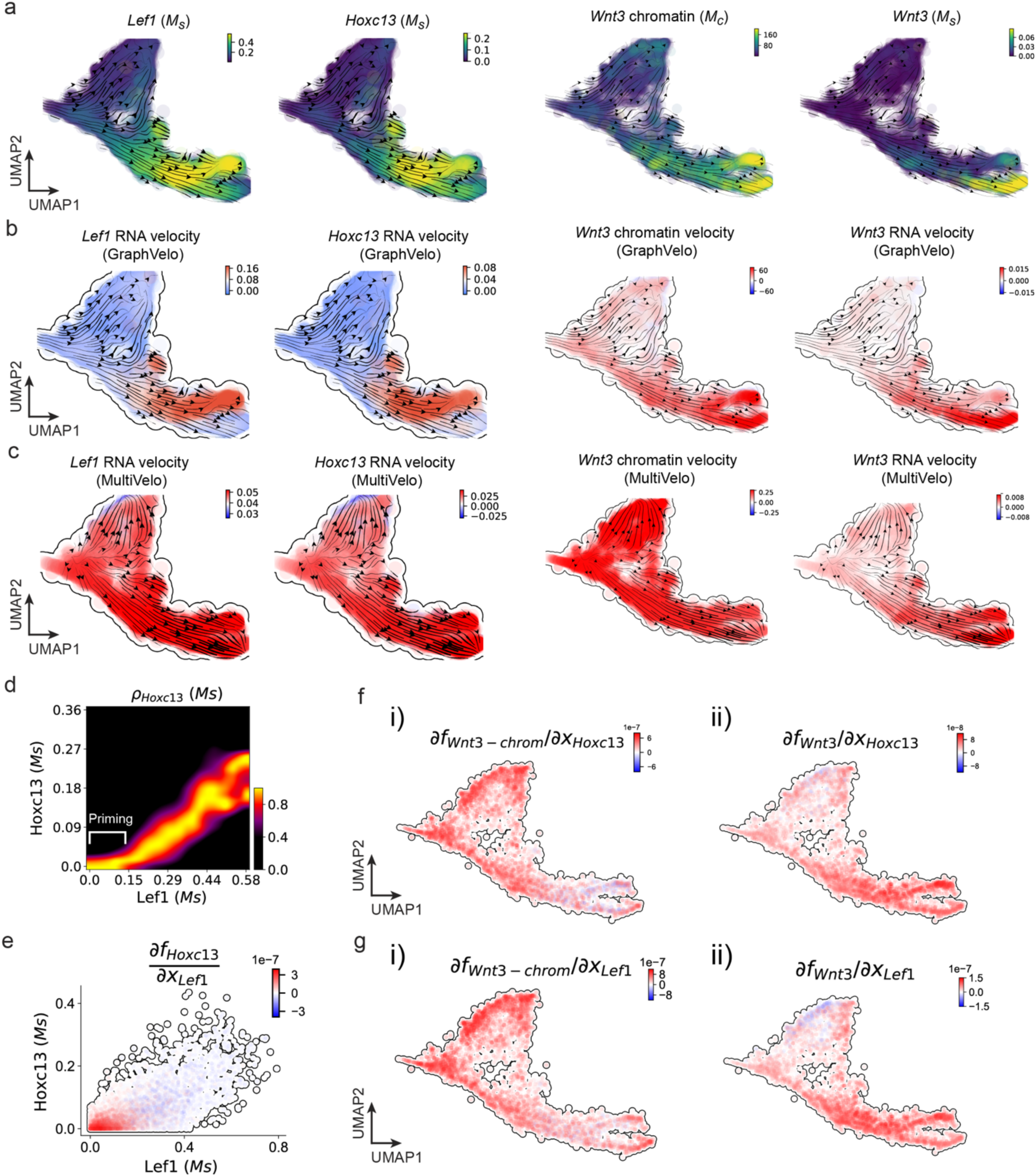
GraphVelo prediction on the regulatory mechanisms of *Lef1-Hoxc13-Wnt3* circuit during mouse skin development. **(a)** Distributions of *Lef1* expression, *Hoxc13* expression, *Wnt3* chromatin openess/accessibility and *Wnt3* expression, respectively, on the projected UMAP vector field. **(b)** Velocities of *Lef1* RNA, *Hoxc13* RNA, *Wnt3* chromatin and *Wnt3* RNA inferred by GraphVelo, visualized on the projected UMAP vector field. **(c)** Velocities of *Lef1* RNA, *Hoxc13* RNA, *Wnt3* chromatin and *Wnt3* RNA inferred by MultiVelo, visualized on the projected UMAP vector field. **(d)** Expression of *Hoxc13* gene versus *Lef1* expression. *Lef1* expression primes the activation of *Hoxc13*. **(e)** Cell-wise Jacobian analyses of *Lef1-Hoxc13* activation cascade. **(f)** Jacobian analyses of regulatory interactions between potential PTF *Hoxc13* and i) *Wnt3* chromatin accessibility or ii) transcription.. **(g)** Jacobian analyses of regulatory interactions between PTF *Lef1* and i) *Wnt3* chromatin accessibility or ii) transcription.

**Extended Data Fig. 20.**
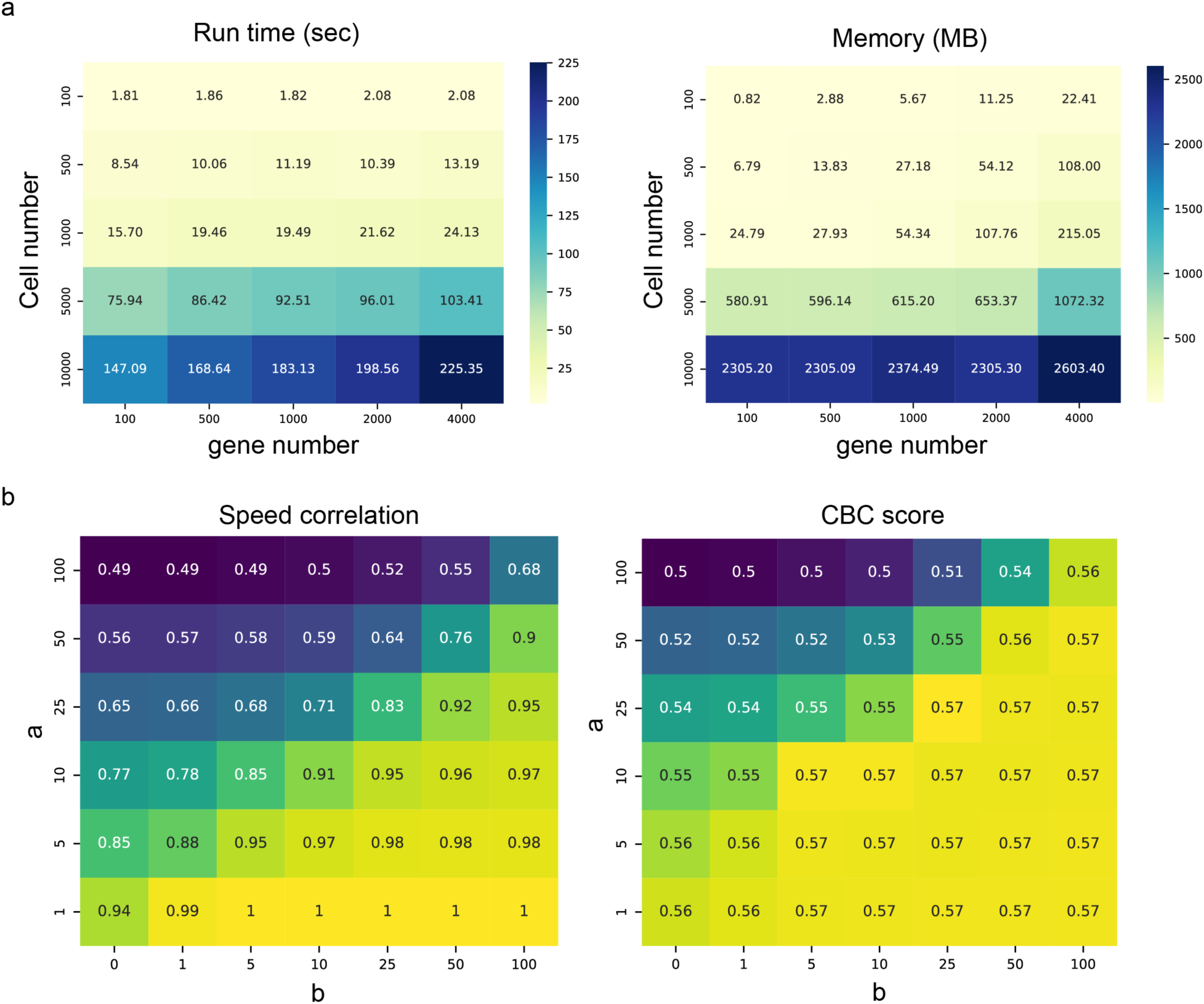
Sensitivity analysis of GraphVelo on simulated and mouse erythroid dataset. **(a)** Runtime (left) and peak memory (right) consumption of GraphVelo with varying numbers of cells and genes on simulated data. **(b)** The speed correlation (left) against GraphVelo result reported in Figure. 3 and CBC score (right) with varying hyperparameters *a* and *b*.

