## Supplemental text for "GraphVelo allows for accurate inference of multimodal velocities and molecular mechanisms for single cells"

### I. Connection of the projection to local linear embedding (LLE)

In this work, we use the neighborhood of a data point to approximate the corresponding tangent space of the data manifold. Such approximation has been used in other applications such as the local linear embedding (LLE)[1]. Here we provide a derivation of representing velocity vectors based on LLE.

Consider a data point $\mathbf{x}_i$ and its neighbors. Applying LLE one has $\sum_{j\neq i} w_{ij}\left(x_j - x_i\right) = \sum_{j\neq i} w_{ij}\delta_{ij} = 0$, with $\sum_j w_{ij} = 1$. The LLE algorithm gives $w_{ij} = \frac{\sum_k C_{jk}^{-1}}{\sum_{lm} C_{lm}^{-1}}$, with $C_{ij} = \left(x_i - x_j\right)\cdot(x_i - x_k)$. Now consider after $\delta t \to 0$ $\mathbf{x}_i$ moves to $\mathbf{x}_i' = \mathbf{x}_i + \boldsymbol{v}_{pi}\delta t$, which should be also within the local linear embedding space. Again with LLE one has $\sum_{j\neq i} w_{ij}'\left(x_j - x_i'\right) = 0$, with $\sum_j w_{ij}' = 1$. Then one has $\boldsymbol{v}_{pi}$=$\sum_{j\neq i}\frac{(w_{ij}' - w_{ij})}{\delta t}\delta_{ij}$ $=\sum_{j\neq i}\phi_{ij}\delta_{ij}$ , with a form similar to that in eq. 1 of the main text.

The above derivation suggests an iterative trajectory propagation algorithm for obtaining the velocities in the new representation:

1) Use a trial $\boldsymbol{v}_{pi}$ to propagate the state and obtain $(\mathbf{x}_i, \mathbf{x'}_i)$. One can propagate backward and forward by $\delta t/2$ and use central difference to estimate the velocity vectors.
2) Apply LLE to get $w_{ij}$ and $w'_{ij}$. I
3) Obtain a new $\boldsymbol{v}_{pi}$
4) Go back to step 1 until the result converges below a threshold.

We numerically tested the above algorithm, the one with the loss function given in eq. 2 of the main text with the parameter $b = 0$ and $b \neq 0$. All the three algorithms worked well on synthetic data. The first two actually outperformed the third one (used in the main text). However, the first two sometime showed numerical instability once applied to real single cell data. Therefore, in subsequent applications, we used the general loss function form in eq. 2, using the direction information from the second term (with $b \neq 0$) to further regularize the projection.

### II. Numerical issues of using eq. 1 for tangent space projection

The relation in eq. 1 provides an algorithm for projecting a measured $\mathbf{v}$ onto $\mathcal{M}$ by minimizing the following loss function,

$$\mathcal{L}(\boldsymbol{\phi}_i) = \left\|\boldsymbol{v}_i - \boldsymbol{v}_{\parallel i}\right\|^2 + \lambda\|\boldsymbol{\phi}_i\|^2,$$

which unfortunately is numerically unstable. The redundancy of the basis vectors leads to coefficients that are not uniquely determined, and failure of the projection. To see the latter, notice that in real data the subspace formed by the displacement vectors only approximates the tangent space $T\mathcal{M}$ locally, and it likely contains small components in the orthogonal space. Then the projection procedure tries to express both $\mathbf{v}_{\parallel}(\mathbf{x})$ and all or part of the remaining $(\mathbf{v}_i - \mathbf{v}_{\parallel})$ as a linear combination of $\boldsymbol{\delta}_{ij}$ with some $|\phi_{ij}| \gg 0$, which can only be counterbalanced by a very large value of $\lambda$. The latter would impose large weight on the regularization. While further data preprocessing may be developed in the future to alleviate the problem, in this work we provided a strategy of adding an additional cosine kernel term for the direction information.

We also experimented with the idea of removing the redundancy through dimension reduction with PCA. The numerical results were not satisfactory, since it converged slowly with the sampling size, which becomes impractical.

### III. Reformulation of Cosine kernel used in the literature in the context of tangent space projection

In the original RNA velocity study, a Cosine kernel has been proposed to transform the RNA velocity vectors between different representations[2]. We re-casted the cosine kernel in the context of the tangent space projection with a form similar to that of eq. 1 in the main text:

$$\mathbf{v}_{\parallel}^{corr}(\mathbf{x}_i) = \sum_{j \in \mathcal{N}_i} \phi_{ij}^{\mathrm{corr}} \boldsymbol{\delta}_{ij},$$

where $\phi_{ij}^{\mathrm{corr}} = P_{ij} - \frac{1}{k}$, and $P_{ij} = \frac{\exp[\cos(\mathbf{v}_i, \boldsymbol{\delta}_{ij})/\sigma]}{\sum_m \exp[\cos(\mathbf{v}_i, \boldsymbol{\delta}_{im})/\sigma]}$ defined in the form of softmax functions, with $\cos(\cdot,\cdot)$ denoting the cosine similarity between two input vectors, $\sigma$ an arbitrary bandwidth parameter, and $k$ the number of neighbors for each cell. Here $P_{ij}$ gives a heuristic transition probability from cell $i$ to $j$. The term $(-1/k)$, called the "density correction", is designed to correct the potential sampling bias where the embedded velocity vectors tend to point towards the direction of regions with high cell density. Li et al. showed that mathematically the cosine kernel asymptotically converges to the correct direction of a velocity vector[3]. However, the correlation kernel loses information about the magnitude of the velocity vectors $\mathbf{v}_i$ (i.e., the *speed*), due to the normalization in the correlation functions. Intuitively, the correlation kernel is qualitatively guided by the physical intuition that a cell has a high tendency to move along the direction of its velocity vector. Here, we used the direction information of $\phi_{ij}^{\mathrm{corr}}$ to help on constraining $\boldsymbol{\phi}_i$.

### IV. Further discussions on applications of GraphVelo to manifolds formed by multi-modal single cell data.

GraphVelo is based on the following mathematical assumption: the manifolds of a given system embedded in two different spaces are homeomorphic so one can establish a one-to-one mapping between the two. Here we use one simple example of scRNAseq/scATACseq multiomic data to illustrate that one can still apply GraphVelo if this assumption is violated. Consider that a cellular system switches between two discrete epigenetic states, and for each epigenetic state there is a corresponding (quasi)continuous transcriptomic manifold. Label the two disjoint manifolds as $\mathscr{M}_1(\mathbf{x})$ and $\mathscr{M}_2(\mathbf{x})$, where $\mathbf{x}$ represents the transcriptomic state. Consider two cells having the same $\mathbf{x}$ but different chromatin state, $c_1$ on $\mathscr{M}_1$ and $c_2$ on $\mathscr{M}_2$. Multiomic data allows distinction between two cells. That is, the neighborhood of $c_1$ is composed of cells on $\mathscr{M}_1$, and the neighborhood of $c_2$ is composed of cells on $\mathscr{M}_2$. Therefore, GraphVelo analyses treat cells on $\mathscr{M}_1$ and $\mathscr{M}_2$ separately.

### V. Feature selection for robustly estimated velocity genes.

In datasets featuring non-differentiating cell types organized in a hierarchical manner, application of RNA velocity may show different kinetic regimes. Analyzing cell fate transition using unreliable velocity genes can lead to emergence of spurious cell state transitions. Different packages subset the well-estimated kinetic genes for further analyses in different scenarios. In the implementation of scVelo, one first regresses out the uncertain genes based on the r-squared coefficient in the steady-state model and further selects trustful velocity genes according to the log-likelihood in the dynamical mode. PhyloVelo relies on monotonically expressed genes to infer transcriptomic velocity, which is a more stringent requirement. However, the monotonic assumptions are counterintuitive and oversimplify the gene expression kinetics along differentiation or disease trajectories due to the non-sequential nature of these cascades. Dynamo provides a biological prior-guided method and filters out the unreliable genes based on the knowledge of lineage information.
